# The role of behavioural flexibility in primate diversification

**DOI:** 10.1101/2020.10.15.341859

**Authors:** Maria J.A. Creighton, Dan A. Greenberg, Simon M. Reader, Arne Ø. Mooers

## Abstract

Identifying the factors that influence species diversification is fundamental to our understanding of the evolutionary processes underlying extant biodiversity. Behavioural innovation, coupled with the social transmission of new behaviours, has been proposed to increase rates of evolutionary diversification, as novel behaviours expose populations to new selective regimes. Thus, it is believed that behavioural flexibility may play an important role in driving evolutionary diversification across animals. We test this hypothesis within the primates, a taxonomic group with considerable among-lineage variation in both species diversity and behavioural flexibility. We employ a time cut-off in our phylogeny to help account for biases associated with recent taxonomic reclassifications and compare three alternative measures of diversification rate that consider different phylogenetic depths. We find that the presence of behavioural innovation and social learning are positively correlated with diversification rates among primate genera, but not at shallower phylogenetic depths. Given that we find stronger associations when examining older rather than more recent diversification events, we suggest that extinction resistance, as opposed to speciation, may be an important mechanism linking behavioural flexibility and primate diversification. Our results contrast with work linking behavioural flexibility with diversification of birds at various phylogenetic depths. We offer a possible dispersal-mediated explanation for these conflicting patterns, such that the influence behavioural flexibility plays in dictating evolutionary trajectories differs across clades. Our results suggest that behavioural flexibility may act through several different pathways to shape the evolutionary trajectories of lineages.

---

Extant species diversity is remarkably variable across the Tree of Life (Willis, 1922; Williams, 1964). For clades of the same age, differences in net diversification rate (i.e. speciation rate minus extinction rate) ultimately drive differences in clade size. Both the external environment (e.g. Badgley, 2010; Kozak & Wiens, 2010; Frey, 2010) and intrinsic features of a lineage (e.g. Raikow, 1986; Heard & Hauser, 1995) can influence net diversification. Despite ongoing study, there remains considerable debate over the factors that lead to differences in diversification rate among lineages, and their relative importance (e.g. Lewontin, 1983; West-Eberhard, 1989; Isaac, et al., 2005; Rabosky, 2009; see review by Wiens, 2017).

Plasticity has been repeatedly proposed to play a major role in shaping evolutionary trajectories in general, and speciation in particular (Baldwin, 1902; Lewontin, 1983; Bateson, 1988; Wcislo, 1989; Odling-Smee, et al., 2003; West-Eberhard, 2003; Pelletier, et al., 2009; also see review in Pfennig, et al., 2010), and theoretical modelling supports its potential influence (e.g. Hinton & Nowlan, 1987; Anderson, 1995; Behera & Nanjundiah, 1995; Ancel, 1999; 2000). Behavioural development and expression often allow for more rapid responses than other forms of plasticity such as induced morphological changes (Duckworth, 2009; Snell-Rood, 2013; West-Eberhard, 2003). Thus, plasticity of behaviour has been hypothesized to play a particularly important role in influencing evolutionary trajectories of animal lineages (Wyles, et al., 1983; Wilson, 1985; West-Eberhard, 2003). The propensity to adopt new behaviours can greatly and quickly alter ecological niches, exposing populations to new selective regimes. This can lead to increased trait disparity among populations (the “behavioural drive hypothesis”; Wyles, et al., 1983; Wilson, 1985); importantly, divergent selection can also lead to eventual speciation. As a result, it has been suggested that behaviourally flexible taxa (i.e. those taxa exhibiting high propensities for behavioural change due to, for example, learning or readiness to transition to new conditions; Sol & Lefebvre, 2000; Audet & Lefebvre, 2017) may undergo faster rates of speciation, and thus overall net rates of evolutionary diversification, compared to less flexible taxa (Sol, et al., 2005; Grant & Grant, 2008; Price, 2008; Sol & Price, 2008; Tebbich, et al., 2010). However, despite support from theoretical modelling (e.g. Price, et al., 2003; Lachlan & Servedio, 2004; Lapiedra, et al., 2013), the idea that behavioural flexibility enhances diversification rates remains contested. Some dispute the extent to which behaviour plays an active role in dictating animal diversity (e.g. Scott-Phillips, et al., 2014). Meanwhile, an alternative hypothesis posits that behavioural flexibility may inhibit, rather than enhance, species diversification: populations that can utilize new resources or transition to new environments are shielding their genomes from bouts of strong directional selection (Bogert, 1949; Huey, et al., 2003).

Duckworth (2009) suggests that behavioural flexibility could both dampen and promote evolutionary rates depending on the time scale. Under this proposed framework, behavioural flexibility inhibits evolution on short time scales by buffering against abrupt environmental changes that may otherwise result in a population bottleneck or strong bouts of directional selection. Over longer time scales, the same behavioural shift can lead to speciation, either by setting the stage for allopatric speciation or by exposing the newly situated population to novel selective regimes (Huey, et al., 2003; Losos, et al., 2004; Duckworth, 2009; see example in Muñoz & Losos, 2018).

Previous studies have provided support for behavioural flexibility both driving (e.g. Yeh, 2004; Yeh & Price, 2004; Tebbich, et al., 2010, Riesch, et al., 2012; Foote, et al., 2016) and inhibiting (e.g. Losos, et al., 2004; Weber, et al., 2004; Shultz, et al., 2005; Gonzalez-Voyer, et al., 2016) evolution. However, many of these studies have primarily considered the effects of behavioural flexibility on microevolutionary change at short time scales (e.g. recent speciation events or population decline). One notable exception is in birds, where behavioural flexibility has been associated with multiple estimates of macroevolutionary lineage diversification (Nicolakakis, et al., 2003; Sol, et al., 2005 – also see Sol, 2003; Sayol, et al., 2019). These studies have employed two proposed correlates of behavioural flexibility: brain size relative to body size (a structural measure presumed and shown elsewhere to be associated with behavioural flexibility; e.g. Lefebvre, et al., 2004) and innovation rate (a behavioural measure) taken from literature surveys. Both large relative brain size and high innovation rates were associated with heightened diversification in birds (Nicolakakis, et al., 2003; Sol, et al., 2005; Sayol, et al., 2019), consistent with the idea that behavioural flexibility positively impacts diversification. Importantly, such tests have yet to be applied across other taxa, making it difficult to generalize the role of behaviour in shaping the Tree of Life.

Here, we explore the relationship between four proxies of behavioural flexibility and several measures of diversification rate in primates, a taxonomic group with considerable among-lineage variation in both behavioural flexibility (Reader & Laland, 2002; Reader, et al., 2011) and extant species diversity (Upham, et al., 2019a; 2019b). Variables associated with diversity of other taxa (e.g. geographic range size and latitude) have been shown to be poor predictors of primate diversification (Arbour & Santana, 2017; Upham, et al., 2019a), leaving a great deal of what shapes extant primate diversity unexplained. We focus on two behavioural measures of behavioural flexibility, the presence or absence of published reports of innovation and of social learning, and two brain size measures widely thought to be associated with ability to exhibit flexible behaviours. Consistent with what has been reported for birds (Nicolakakis, et al., 2003; Sol, et al., 2005; Sayol, et al., 2019) and Duckworth (2009)’s proposal that behavioural flexibility promotes diversification events over longer evolutionary time scales, we predict that our separate measures of behavioural flexibility will covary positively with diversification rates across primate lineages. To examine how this association changes at different phylogenetic depths, we examine how behavioural flexibility correlates with diversification over both shallower and deeper time depths in our phylogeny. Understanding how behaviour and ecology may interact to shape evolutionary patterns provides a glimpse into some of the processes that have shaped primate biological diversity and could, in turn, dictate future diversity.

## METHODS

### Data

#### Diversification Rate

Estimating diversification rates is challenging because it depends on an accurate assessment of both the taxonomic richness and divergence time of a given lineage. A previous study testing whether behavioural flexibility drives shallow divergences used subspecies per species as a measure of subspecific diversification in birds (Sol, et al., 2005). Using this subspecies metric could introduce considerable bias when applied in primates, however, as primate taxonomic richness has changed drastically over recent decades (Tattersall, 2007; Groves, 2014; Rylands & Mittermeier, 2014), with much of this change attributed to application of the ‘phylogenetic species concept’ (PSC) and its tendency to raise former subspecies and variants to the full species rank (Tattersall, 2007). Importantly, it has been suggested that these elevations in subspecies status have been biased toward certain taxa (Isaac, et al., 2004), which would lead to inconsistent estimates of species versus subspecies richness across lineages. Studies using other estimates of primate diversification (i.e. diversification analyses using TreePar; Stadler, 2011) have been hindered by the applications of the PSC, particularly when it comes to accurately estimating shallow divergences (Springer, et al., 2012). Modern primate phylogenies are also not reflective of modern primate taxonomies, as they include some phylogenetic species and omit others, preventing us from using recently described evolutionary rate measures (see, e.g., Jetz, et al., 2012; Mitchell & Rabosky, 2017) that rely on a comprehensive phylogeny with consistent species designations among clades. Instead, we used well-resolved “lineages” that putatively reflect stable species complexes. We started with the most widely used, dated primate tree publicly available at the time of this study, the GenBank taxonomy consensus tree provided on the 10kTrees website (version 3) (Arnold, et al., 2010), containing 301 tips. We then created a time cut-off in the tree at the time when we determined a majority of robust biological species described in Honacki et al. (1982) had evolved (1.1mya). We chose Honacki et al. (1982) as it was the last major primate taxonomy published before the introduction of the PSC (Cracraft, 1983). After creating this time cut-off, we subsequently eliminated shallow divergences occurring after 1.1mya from the consensus tree (see Figure S1). Each branch in the tree that was extant at 1.1mya was retained in the tree and designated as a stable “lineage”. We then additionally pruned species from this phylogeny that were no longer recognized by modern taxonomies (IUCN/SSC Primate Specialist Group, 2018). This resulted in 241 identifiable lineages to compare in terms of taxonomic richness and divergence times. Using the most recently published primate species and subspecies list from the IUCN/SSC Primate Specialist Group (2018), we referenced taxonomic and phylogenetic works to assign each of the 705 species and subspecies to one of these 241 lineages. This allowed us to assign each lineage an agnostic “taxon richness” score (i.e. the sum of all monotypic species and subspecies in a lineage; see Figure S1) that accounts for the discrepancies in subspecies elevations across lineages. Each species or subspecies listed by the IUCN/SSC Primate Specialist Group (2018) was also assigned to one of the 66 genera named in our 10kTrees phylogeny (Arnold, et al., 2010). After eliminating two individual species and one genus in our tree that could not be resolved using these methods (see supplementary material), our study considered 703 taxa (species or subspecies) assigned to 239 lineages and 65 genera.

To estimate lineage diversification rate we used the method-of-moments estimator (Magallon & Sanderson, 2001) that divides the natural log of “taxon richness” (species and subspecies) by lineage stem age to produce a ‘Taxa per Lineage Diversification Rate’ that should be less biased by recent subspecies elevations. This method was repeated at the genus-level where the natural log of “taxon richness” for each genus was divided by the stem age of that genus (hereafter ‘Taxa per Genus Diversification Rate’). We note that all log transformations referenced hereafter refer to natural log transformations (loge). Lastly, we created a second, and perhaps more conservative estimate of genus diversification rate where richness scores were generated using the number of lineages (n=239) per genus, rather than the number of taxa described by the IUCN/SSC Primate Specialist Group (2018), hereafter ‘Lineage per Genus Diversification Rate’. A few genera were not monophyletic on our tree, and we considered these on a case-by-case basis ultimately removing five from our genus-level analysis (see the supplementary material).

We illustrate these methods and present associated calculations using genus *Aotus* as an example in the supplementary material to provide some considerations on the potential uses and limitations of this approach.

#### Behavioural Proxies of Behavioural Flexibility

We focused on two key behaviours to infer behavioural flexibility: innovation and social learning. Innovation and social learning are both important in determining the macroevolutionary effects of behavioural flexibility because multiple individuals must acquire an innovation through social learning or independent innovation to have population-level effects (Wyles, et al., 1983; Wilson, 1985; Duckworth, 2009). In addition to facilitating the transmission of innovations throughout a population, social learning can also be a valuable measure of population-level behavioural flexibility on its own as it reflects the ability of individuals within a population to pick up behaviours that are novel to them but not necessarily novel to the population. We note that social learning and innovation are taxonomically widespread (Reader & Biro, 2010; Snell-Rood, et al., 2015). However, we assume that species with no published accounts of social learning or innovation are likely relying on these behaviours infrequently. Moreover, the innovation and social learning data used here have been positively associated with other measures of behavioural flexibility (e.g. brain size measures; Reader, et al., 2011; Navarrete, et al., 2016). For innovation, we focus on technical innovations (classified as those involving tool use following Navarrete, et al., 2016) because these more easily defined behaviours have been linked to complex cognition (Overington, et al., 2009), and reports of other classes of innovation (e.g. food type innovation) can be highly influenced by opportunistic events (Ducatez, et al., 2015). Combined with the fact that taxa with reports of technical innovation also tended to be those with evidence of other innovation types (see data in Navarrete, et al., 2016), this likely makes technical innovation a robust estimate of innovativeness across primates.

Counts of innovation and social learning per lineage came from Reader et al. (2011) and Navarrete et al. (2016). Reader et al. (2011) established this dataset through a survey of over 4000 published articles for examples of social learning and behavioural innovation. Reader et al. (2011, p. 1018) define an innovation as the tendency to “discover novel solutions to environmental or social problems”. These data were later subdivided into different innovation categories by Navarrete et al. (2016), including ‘technical’ innovations, defined as innovations involving tool use. Reader et al. (2011, p. 1018) define social learning as the tendency to “learn skills and acquire information from others”, including instances of social learning from both kin and unrelated individuals. Social learning was often inferred from observational data in the original reports.

As an alternative to treating behavioural data as a binary metric (e.g. presence or absence of innovativeness in Ducatez, et al., 2020) some studies have used “rate” measures of behaviours: residuals from a log-log plot of the total number of recorded instances of a behaviour (e.g. social learning) and an estimate of research effort (e.g. the number of papers published on that taxa; e.g. Sol, et al., 2005; Reader, et al., 2011; Navarrete, et al., 2016; Ducatez, et al., 2020). However, the choice of how to model the relationship between total number of recorded innovation or social learning instances and research effort matters considerably when creating these residual rate measures. For our data, the relationship between the total number of innovation or social learning instances and research effort was non-linear. This made the choice of model structure non-trivial, with different models proving difficult to justify over one another. Residuals from these models additionally showed further structural issues, including failure to meet assumptions about homoscedasticity. Thus, we opted to use binary measures, which allowed us to use data imputation methods to minimize biases caused by under-studied taxa, statistically account for potential biases associated with summarizing behavioural data at higher taxonomic levels, and run simulations to address the assumptions underlying our analyses. We therefore assigned each lineage or genus binary scores of 1 (presence of innovation or social learning) or 0 (absence of innovation or social learning). Further considerations regarding the use of literature-based estimates of behavioural flexibility can be found in the supplementary material.

#### Structural Proxies of Behavioural Flexibility

Literature-based evidence for behavioural flexibility across taxa has its limitations and so we chose to also consider structural correlates of behavioural flexibility. It is widely thought that particular brain regions are associated with flexible behaviour – particularly the neocortex (see, e.g., Keverne, et al., 1996; Mikhalevich, et al., 2017) and cerebellum (Vandervert, 2003; Vandervert, et al., 2007; Barton, 2012). Therefore, in addition to total brain size (relative to body mass), we considered the sum of neocortical and cerebellar volumes relative to rest of total brain volume as another proxy for behavioural flexibility. Using both behavioural measures and structural correlates of behavioural flexibility, we were able to compare the consistency of results across different proxies for behavioural flexibility.

Lineage-level estimates for all brain measures were calculated by taking the geometric mean of taxon volumes for each lineage. Brain volume relative to body size (hereafter ‘relative brain volume’) was estimated by regressing lineage-level estimates of logarithmic endocranial volume, in cm^3^, (ECV) (Powell, et al., 2017) (hereafter ‘brain volume’) as a function of logarithmic body mass, in grams (Jones, et al., 2009), and retaining the residuals (Dunbar & Schultz, 2007). Notably, previous studies (e.g. Sayol, et al., 2019) have used phylogenetically corrected residuals to account for an effect of body size when using similar brain size measures. However, for our data, residuals obtained from phylogenetic models were skewed such that lineages with larger body mass consistently had larger residual brain volumes (see Figure S2), and thus we opted to use ordinary least square residuals that were more orthogonal to lineage body size. Consistent with relative brain volume, neocortex and cerebellum volume relative to rest-of-brain volume (hereafter ‘relative neocortex and cerebellum volume’) was estimated by taking the residuals from a log-log regression of the combined neocortex and cerebellum volumes on the rest-of-brain volumes (i.e. total brain volume minus neocortex and cerebellum volumes) taken from Navarrete et al. (2018) and the compilation in DeCasien & Higham (2019). Genus-level estimates for structural proxies of flexibility were calculated by taking the mean of the lineage estimates within each genus. Further details, and considerations regarding the use of structural proxies of behavioural flexibility, and residual brain measures can be found in the supplementary materials along with correlation coefficients for all predictor variables (Table S1).

Phylogenetic signal of all predictor and response variables are reported in the supplementary material (Table S2).

### Analysis

All analyses used R version 3.6.3 (R Core Team, 2020).

#### Trait Imputation

While we collected data for our predictor variables from the most comprehensive datasets and compilations available, research biases and the persistent reassessment of primate taxonomy has resulted in inconsistent data coverage across lineages, and there were still many lineages that were missing data (see Figure S3). To maximize our evolutionary inferences on diversification and allow for the inclusion of data-poor lineages, we chose to impute missing predictor variables using phylogenetic imputation methods (see supplementary material for details and reports of predictive accuracy from model cross-validation; Table S3 and Figure S4). Data on relative neocortex and cerebellum volume were sparse and unevenly distributed across the phylogeny (82.4% of lineages missing data; Table S3), making it infeasible to reliably impute missing values. We thus only ran models of relative neocortex and cerebellum volume on the original, non-imputed dataset. All of the regressions we report below were repeated for the original, non-imputed datasets (see Results and the supplementary material Tables S4 to S6) and except as noted gave similar results.

#### Predictors of Diversification

To assess the relationship between our measures of behavioural flexibility and diversification rate at the lineage-level (Taxa per Lineage Diversification Rate) we used phylogenetic generalized least squares (PGLS) regressions implemented using the “gls” function in the nlme package (Pinheiro, et al., 2020), including the 10kTrees consensus tree for our 239 defined lineages as the phylogenetic backbone. PGLS is a common regression method used to investigate evolutionary associations while accounting for the fact that closely-related lineages tend to be similar (e.g. in body size, life history and ecology; see Freckleton, et al., 2002). Continuous data were log-transformed, centred with respect to the mean, and scaled by 2 standard deviations in all models for both the lineage and genus-level analyses to make effect sizes comparable to those reported for binary variables (Gelman, 2008). After imputing missing values, our dataset contained 54 lineages scored as having evidence of social learning (scored as 1) and 28 lineages scored as having evidence of technical innovation (scored as 1). Wyles et al. (1983) predicted accelerated evolution in species with a dual capacity for innovation and social propagation of new behaviours, therefore, we also tested a combined measure of technical innovation and social learning. In this combined measure 26 lineages with the presence of both behaviours were scored as 1, and those exhibiting only one or neither behaviour were scored as 0.

To assess the relationship between our measures of behavioural flexibility and diversification rate deeper in the tree we repeated the same analysis at the genus-level using two different estimates of diversification rate: Taxa per Genus Diversification Rate and the more conservative Lineage per Genus Diversification Rate. After imputing missing values, our dataset of 60 genera contained 21 genera scored as having evidence of social learning, 9 genera scored as having evidence of technical innovation and 8 genera with evidence of both behaviours.

Body mass and attendant life history traits have been predicted to impact diversification rates in some taxa, albeit with conflicting results (see, e.g., Cardillo, et al., 2003; Paradis, 2005; Fontanillas, et al., 2007; Thomas, et al., 2010), and body mass is closely correlated with many primate life history traits (e.g. age at first reproduction, maximum lifespan; Charnov & Berrigan, 1993; Purvis, et al., 2003; Street, et al., 2017). To examine whether our results could stem from confounding effects of body mass and its correlates, we ran PGLS analyses to test body mass as an independent predictor of our diversification rate measures. Results from these tests were non-significant across all measures of diversification (see supplementary material Tables S4 to S6).

We re-ran all PGLS analyses reported above after eliminating the great apes, which are characterized by slow life histories, which may counteract the evolutionary effects of behavioural flexibility by slowing their evolutionary response time (Price, et al., 2003). For tests that proved significant, we then reran the same test 100 times over a sample of 1000 trees from the tree block available in 10kTrees (Arnold, et al., 2010) to confirm results were consistent after accounting for phylogenetic uncertainty.

#### Genus-Level Simulations

Genera were considered behaviourally flexible if any of their daughter lineages had evidence of innovation or social learning, which potentially introduces a statistical bias if more lineage-rich genera (which will generally have higher diversification rates) are more likely by chance to include at least one lineage that expresses technical innovation or social learning. To account for this possible bias, we simulated the neutral evolution of technical innovation and social learning across the primate phylogeny 1000 times using the symmetrical rate Mk model of discrete trait evolution (Lewis, 2001). We opted to model the evolution of these behaviours under a symmetrical model of trait evolution since with smaller datasets such as ours, there is little power to prefer asymmetric models (Mooers & Schluter, 1999). From these stochastic distributions of the two traits, we repeated our analyses of diversification rate and created a distribution of expected effect sizes under a null evolutionary scenario. More diverse genera may also be more likely, by chance, to have well-studied lineages, which could create a bias toward observing the presence of innovation or social learning in diverse genera (Ducatez & Lefebvre, 2014). Although data did not suggest that lineage-rich clades were more likely to have intensely investigated lineages in our dataset (see Figures S5 and S6), we nonetheless opted to take research effort into account in our analysis and expanded upon our first simulation to consider a scenario where the presence of technical innovation or social learning may go unobserved if insufficient research effort was directed at a lineage. We took the 1000 simulations of neutral evolution of technical innovation and social learning, and then randomly ‘evolved’ research effort for each simulation, independently of the evolution of technical innovation and social learning. We then converted the presence of these behaviours to absences in our simulated datasets if a lineage’s ‘evolved’ research effort was below the minimum threshold of studies for a lineage with observed technical innovation (as it contained the higher research effort threshold than social learning; Figure S7). By repeating our analyses of diversification rate with these new simulated datasets where lineages with low research effort were assigned ‘hidden states’, we were able to see if a bias towards having better studied lineages in diverse genera could drive a positive association between behavioural proxies of flexibility and diversification rate by chance and independent of biological mechanisms. Importantly, this simulation makes the assumption that research effort and behavioural flexibility are independent; if this is not true such that behaviourally flexible lineages attract research effort (see discussion in the ‘Research Effort Bias’ section of the supplementary material) then the results of this simulation would actually be conservative. We detail both simulations, along with considerations about using research effort as a covariate in binary models (Ducatez, et al., 2020), in the supplementary material.

### Ethical Note

This research was comparative and was based on data available in previously published literature.

## RESULTS

### Lineage-Level Predictors of Diversification

We found no support for an association between behavioural flexibility and diversification rates when testing our measures of behavioural flexibility at the lineage-level (results summarized in Figures 1 and S8; Table S4). Social learning (p= 0.171), technical innovation (p= 0.792), the combined presence of technical innovation and social learning (p= 0.979), relative brain volume (p= 0.215), relative neocortex and cerebellum volume (p= 0.664), and body mass (p= 0.764; Table S4) did not exhibit noteworthy associations with Taxa per Lineage Diversification Rate in either direction.

**Figure 1:**
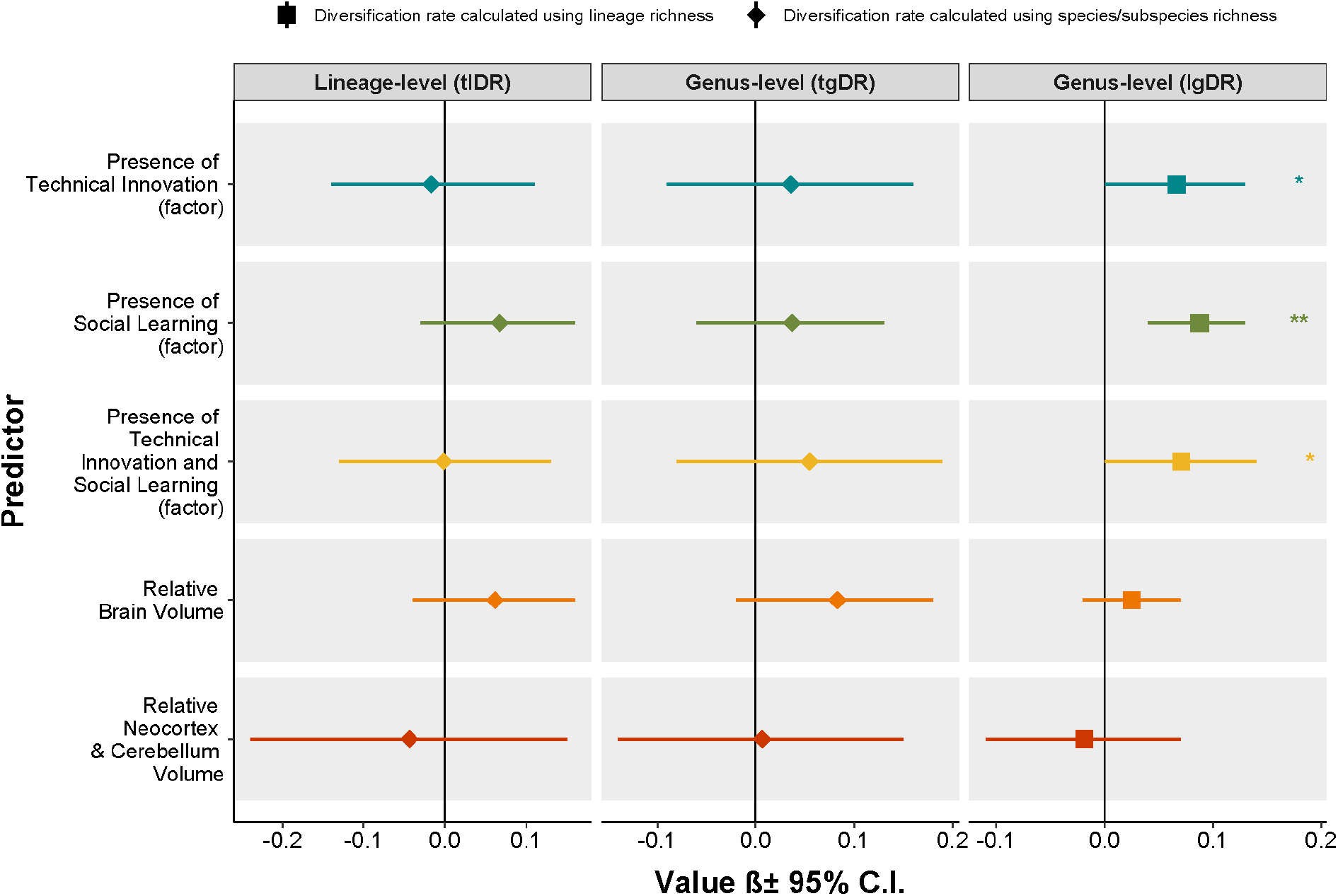
Results from PGLS analyses showing the association of proxies of behavioural flexibility with Taxa per Lineage Diversification Rate (DR), Taxa per Genus Diversification Rate and Lineage per Genus Diversification Rate across primates. 95% confidence intervals are represented by horizontal lines around the associated value. Diamonds indicate diversification rates estimated with species/subspecies richness, and squares indicate diversification rates estimated with lineage richness. Results presented include imputed data when available and brain measures were standardized (ln x/(2sd)). Significance indicated as: +P≤0.1; *P≤0.05; **P≤0.01.

### Genus-Level Predictors of Diversification

Technical innovation (p= 0.577), social learning (p= 0.442), the combined presence of technical innovation and social learning (p= 0.411), relative neocortex and cerebellum volume (p= 0.930), and body mass (p= 0.204) did not exhibit noteworthy associations with Taxa per Genus Diversification Rate (Figures 1 and S8; Table S5). Relative brain volume was insignificantly (p= 0.108), but positively, related to diversification rate, and this relationship became significant when we considered the non-imputed dataset (p= 0.004).

In contrast to the other diversification measures, Lineage per Genus Diversification Rate was positively associated with all three behavioural measures of behavioural flexibility (results summarized in Figures 1 and S8; Table S6). Genera with records of technical innovation were shown to have a faster mean Lineage per Genus Diversification Rate (0.136 lineages my^−1^) than those with no reports (0.070 lineages my^−1^; ß [95% CI] =0.066 [0.002-0.130]; p= 0.048). Genera with records of social learning similarly exhibited a faster mean Lineage per Genus Diversification Rate (0.137 lineages my^−1^) than those without (0.049 lineages my^−1^; ß [95% CI] =0.088 [0.043-0.132]; p< 0.001). Genera with reports of both technical innovation and social learning also exhibited a faster mean Lineage per Genus Diversification Rate (0.153 lineages my^−1^) compared to those with evidence for only one or neither behaviour (0.068 lineages my^−1^; ß [95% CI] =0.085 [0.018-0.151]; p= 0.015). These results were consistent with results run over a sample of 1000 trees from the posterior distribution, i.e. “tree block”, available from 10kTrees (Arnold, et al., 2010), indicating results were not biased by our choice of phylogeny (Figure S9). Based on our simulation testing for a lineage-richness sampling bias, we found that these effect sizes were unlikely to be due to chance alone for technical innovation (p= 0.028), social learning (p= 0.002) and the combined presence of both behaviours (p= 0.008) (see Figure S10 and supplementary results). After expanding our simulation to consider a research effort bias, significance of these effects remained for social learning (p= 0.019) and its combined presence with technical innovation (p= 0.027), but the effect of technical innovation alone was no longer nominally significant (p= 0.109) (see Figure S11 and supplementary results). No structural proxies of behavioural flexibility shared notable associations with Lineage per Genus Diversification Rate (Figures 1 and S8; Table S6).

Results of all PGLS analyses reported above were qualitatively consistent after removing great apes (see supplementary material Tables S7 to S9).

## DISCUSSION

We find little to no compelling support for an association between our proxy measures of behavioural flexibility and recent primate diversification rates of young species and subspecies, however, we do find some evidence supporting a positive association when looking at the diversification of older primate lineages. This pattern could be explained in several ways, and interpretation of these results depend on the assumptions one makes about our different measures of diversification rate.

One benefit of our study design is that it allowed us to consider taxa that are commonly overlooked (i.e. subspecies and species omitted from higher order phylogenies), many of which likely represent very recent splitting events. The weak associations with behavioural flexibility that we observed at shallower time scales could be explained in several ways. On one hand, this pattern could reflect biases in describing taxa (species and subspecies) among groups. While our time cut-off, and use of both species and subspecies, mitigates against biases associated with the elevation of subspecies under the PSC, if less flexible species are more likely to have a larger number of taxa described overall (e.g. based on regional biases in designating species or subspecies that covary with locations of inflexible clades) then this could obscure underlying biological patterns. If this is the case, then behavioural flexibility may indeed enhance diversification by acting on speciation, and we observe stronger associations when ignoring shallow splits because our diversification rate metric calculated using lineage-richness is less biased by this phenomenon. On the other hand, if we take our results at face value, a pattern of stronger associations deeper in the phylogeny could indicate that time plays an even larger role in the relationship between behavioural flexibility and diversification than previously suggested by Duckworth (2009).

Under Mayr’s ‘ephemeral speciation model’ (Mayr, 1963; Rosenblum, et al., 2012) and the related ‘ephemeral divergence hypothesis’ (Futuyma, 1979; 2010), divergence can occur rapidly and often, but many newly diversifying lineages do not persist, instead being eradicated via extinction or ‘reabsorption’ by hybridization (see, e.g., Rosenblum, et al., 2012). It is possible that a number of the described species and subspecies used here (from the IUCN/SSC Primate Specialist Group, 2018) represent such ephemeral diversification events, especially considering that the PSC has facilitated the splitting of very closely related populations. If so, our results could be explained if behavioural flexibility buffers against extinction through behavioural shifts. Behavioural flexibility would then promote diversification through bolstering lineage persistence rather than the rate of splitting; and would be revealed when comparing the accumulation of lineages that have escaped extinction (i.e. looking deeper in the tree). This would be complementary to findings from Arbour & Santana (2017), who show that decreased extinction preceded a shift to increased evolutionary rates in the most speciose primate family (Cercopithecidae), and with evidence suggesting behavioural flexibility is beneficial for population persistence in birds (e.g. Shultz, et al., 2005; Rossmanith, et al., 2006; Sol, et al., 2007, Ducatez, et al., 2020). Under this scenario, the positive associations we find are not the result of behavioural flexibility enhancing evolutionary rates by facilitating divergence events, and instead indicate that extinction resistance may be an important mechanism linking behavioural flexibility and primate diversification. Our results stand in contrast with consistent reports of behavioural flexibility enhancing diversification of birds even at shallow phylogenetic depths (Nicolakakis, et al., 2003; Sol, et al., 2005; Sayol, et al., 2019).

A potential explanation for differing effects of behavioural flexibility in birds versus primates could be that the heightened dispersal capabilities (i.e. flight) of birds makes behavioural flexibility particularly beneficial for these taxa when it comes to establishing in new environments. This could cause behavioural flexibility to enhance bird speciation via increased success in dispersal events while also buffering against extinction – leading to the strong effects that have been observed on their net rates of evolution (Nicolakakis, et al., 2003; Sol, 2003; Sol, et al., 2005; Sayol, et al., 2019). Comparatively, primates have much more limited dispersal capabilities and thus behavioural flexibility may buffer against extinction while having little effect on promoting allopatric establishment and subsequent speciation. If true, it is possible that behavioural flexibility affects evolutionary trajectories via different mechanisms in these clades.

Directly testing whether behavioural flexibility buffers against extinction may be difficult because measuring extinction rates at macroevolutionary scales is notoriously imprecise (Rabosky, 2010; Louca & Pennell, 2020). Future studies could test proxies of primate behavioural flexibility against estimates of contemporary extinction vulnerability (see, e.g., Nicolakakis, et al., 2003; Ducatez, et al., 2020). Another prediction concerns the rate of genetic divergence within flexible versus less flexible, recent clades (e.g. subspecies within species): our conjecture would be that these would not be different, because flexibility does not lead to increased divergence over the short term.

Notably, not all of our results are consistent with a significantly positive association between behavioural flexibility and diversification of older primate lineages. While the non-significant association between Lineage per Genus Diversification Rate and neocortex and cerebellum volume might be explained by data limitations (discussed below), the non-significant association with relative brain volume illustrates that not all proxies of behavioural flexibility capture the same thing. While relative brain volume is a known structural correlate of measures of primate and avian behavioural flexibility (Lefebvre, et al., 2004), the brain has many functions and thus is a less direct measure of behavioural flexibility than behavioural measures. Measures of brain size have been linked to diversification rate in birds (Nicolakakis, et al., 2003; Sol, et al., 2005; Sayol, et al., 2019), but notably the sample for birds is much larger, and is likely to contain a broader range of variation, which would make it easier to detect a brain size effect. Additionally, results of our simulation accounting for potential research effort biases indicate that a positive association between technical innovation and diversification rate in primate genera could be driven by a research effort bias. However, this simulation assumes that research effort and behavioural flexibility are independent and, if instead research effort tends to be directed toward groups where innovation is expected, then this research bias may actually have a low contribution to the observed relationship between technical innovation and diversification rate. Thus, our results paint a complicated picture, and emphasize the need to compare methodologies and measures, and avoid a focus on single explanations for complex phenomena.

Our results provide some support for primate behavioural flexibility being associated with older as opposed to more recent diversification events, however, there are several caveats on the limits of our inferences. First, one measure thought to be associated with behavioural flexibility – neocortex and cerebellum volume – had very little data available for lineages reported as inflexible (e.g. among lemurs, tarsiers, titis and sakis). This meant our ability to test this measure as a driver of diversification was limited. Additionally, to examine the relationship between different measures of behavioural flexibility and diversification rate, we had to run multiple tests, increasing the likelihood of observing Type I error. Our study is also limited by the lineages present in the phylogeny. Available primate phylogenies, (e.g. Arnold, et al., 2010; Upham, et al., 2019b) do not reflect the most recent taxon lists, and such limited taxonomic scope prevented us from using diversification rate estimates that require a fully-resolved tree (e.g. the DR measures from Jetz, et al., 2012). Lastly, many factors will contribute to true diversification, and although we test two (e.g. body mass and a “great ape effect”; see supplemental results), there are many biological and ecological factors (e.g. geographic discontinuities, habitat affiliations, or presence on islands) yet to be explored that could mediate any relationship between behavioural flexibility and primate diversification rate.

While our results provide some evidence for the hypothesis that behavioural flexibility drives diversification of primate lineages, they raise important questions about its underlying mechanisms. Importantly, our results are not consistent in supporting a significant association between behavioural flexibility and diversification rate, and depend on the proxy measures employed. The positive associations we do find may point to behavioural flexibility dampening extinction of young primate lineages, rather than accelerating diversification via splitting of behaviourally shifted individuals/populations. Thus, expanding tests of whether and how behavioural flexibility is associated with diversification to other taxa may help us interpret the results we report here.

## ACKNOWLEDGEMENTS

This work was supported by the Natural Sciences and Engineering Research Council of Canada (NSERC) Discovery Grants Program (SMR and AOM, grants #2017-04720 and #2019-04950) and the Canada Foundation for Innovation (SMR, grant #29433). MJAC was supported by awards from the Biodiversity, Ecosystem Services, and Sustainability (BESS) program and McGill University. We thank the Reader and Guigueno labs, Brian Leung, and Hans Larsson at McGill, William Wcislo, and the Crawford Lab for Evolutionary Studies at SFU for feedback on study design and interpretation, as well as Isabella Capellini and three anonymous reviewers for comments on previous versions.

Data and code will be made available through Dryad or a similar repository.

