## Supplementary Materials for "The role of behavioural flexibility in primate diversification"

### “The role of behavioural flexibility in primate diversification” — Supplementary Material

SUPPLEMENTARY METHODS:

*Diversification Rate*

When assigning species and subspecies listed by the IUCN/SSC Primate Specialist Group (2018) to each lineage >1.1my old in the 10kTrees consensus phylogeny (Arnold, et al., 2010), there were two species that could not be assigned to any of our lineages, and were removed as a result: *Galagoides kumbirensis* (Svensson, et al., 2017) and *Callithrix humilis* (van Roosmalen, et al., 1998). In addition, one genus, *Presbytis* (comprising the surilis), was so unresolved that lineage richness was impossible to estimate given the lineages available in 10kTrees. This genus and its comprised species were also removed prior to analyses.

We estimated ‘Taxa per Lineage Diversification Rate’ using the method-of-moments estimator based on stem age, i.e. the natural log of richness divided by the stem age of that lineage (see Figure S1) (Magallon & Sanderson, 2001). Notably, all log transformations referenced in this document and in the main text refer to natural log transformations ($\log_e$ or ln). In the example in Figure S1B, Taxa per Lineage Diversification Rate for the *Aotus azarai* lineage would be log(3)/1.29 taxa per my. For genus-level diversification rates we delineated genera in our phylogeny and extracted stem ages for each genus, and subsequently used the method-of-moments approach for two estimates of diversification rate. For the first estimate (hereafter ‘Taxa per Genus Diversification Rate’) richness score was estimated based on assignments of the same 705 taxa method-of-moments estimates. ‘Taxa per Genus Diversification Rate’ thus is based on “taxon richness” (so, for *Aotus* in Figure S1, Taxa per Genus Diversification Rate=log(13)/19.49 taxa per my, where 19.49my is the stem age of genus *Aotus*, and 13 is the number of species and subspecies in the genus). For the second estimate, ‘Lineage per Genus Diversification Rate’, richness was equal to the sum of >1.1 million-year old lineages from our tree (so, for *Aotus* in Figure S1, Lineage per Genus Diversification Rate=log(5)/19.49 lineages per my).

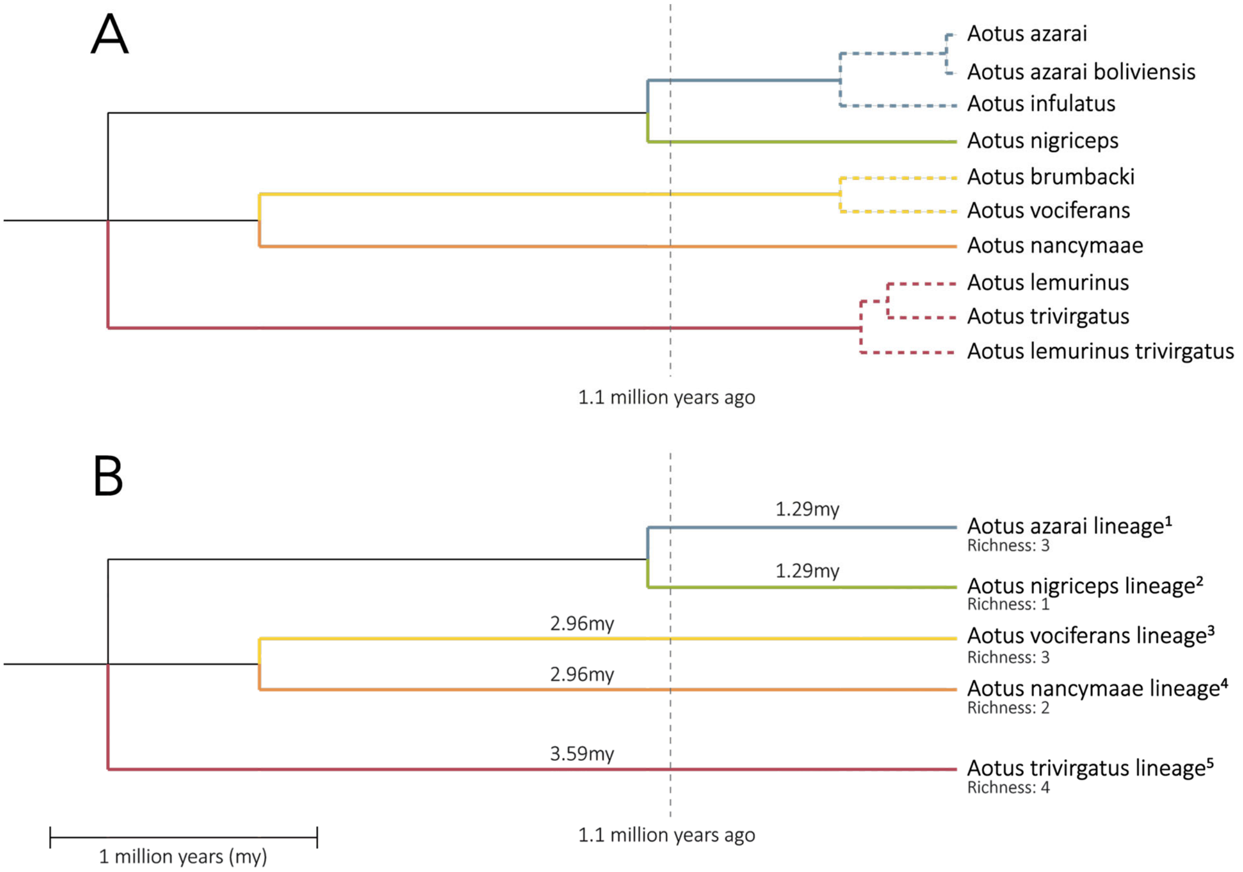


Figure S1: Illustration of the method used to estimate diversification rate at the lineage-level using genus *Aotus* as an example. (A): the 10kTrees consensus phylogeny for *Aotus* where splitting events in the tree occurring more recently than 1.1 million-years ago are represented by dotted lines. (B): the revised phylogeny after collapsing taxa diverging after the 1.1 million-year time cut-off to their corresponding 5 lineages, and appending the taxon richness (total species and subspecies) from the 2018 IUCN taxonomy. Branch lengths (=stem age) are labelled in million years (my). The stem genus age for *Aotus* is 19.49 my (not shown). Species and subspecies listed by the IUCN/SSC Primate Specialist Group (2018) were assigned to lineages to create taxon richness estimates: [1]*Aotus azarae azarae*, *Aotus azarae boliviensis*, *Aotus azarae infulatus*; [2]*Aotus nigriceps*; [3]*Aotus vociferans*, *Aotus brumbacki*, *Aotus jorgehernandezi*; [4]*Aotus nancymaae*, *Aotus miconax*; [5]*Aotus trivirgatus*, *Aotus lemurinus*, *Aotus griseimembra*, *Aotus zonalis* (nomenclature here follows IUCN/SSC Primate Specialist Group, 2018).

We found that there were several instances of non-monophyletic genera within the phylogeny (*Galagoides*, *Otolemur*, *Galago* and *Euoticus*). We elected to retain these lineages in our lineage-level tree (as their unresolved nature may simply be due to taxonomic issues; Masters, et al., 2017), but removed these genera from genus-level analysis as it is unclear how to assign a stem age to these clades. We additionally removed the genus *Semnopithecus*, which comprised a single lineage nested within the genus *Trachypithecus*. Thus, while our lineage-level dataset contained lineages from 65 genera, our genus-level analysis considered only 60 genera. Lineages of *Cercopithecus* were also resolved as polyphyletic in the phylogeny, with a small fraction of lineages forming a clade sister to *Erythrocebus*. Since a majority of *Cercopithecus* lineages were resolved as monophyletic, we opted to assign *Cercopithecus* the divergence date that separates it from the clade of *Chlorocebus* and *Erythrocebus*; we note that removing *Cercopithecus* in its entirety did not qualitatively change the results we report here.

We note that the methods of moments approach to measuring diversification rate does have its limitations, specifically that it can result in a loss of relevant information (i.e. phylogenetic distances between closely related species). This method additionally relies on taxonomic assignments that inherently carry some subjectivity since what is considered a subspecies, species or genus is ultimately determined by taxonomists who must make decisions based on whatever e.g. genetic, morphological, and geographical information is available for a given population. However, this method allowed us to incorporate shallow divergences (in terms of both species and subspecies), which is important since current primate phylogenies do not contain comparable species numbers across clades; applications of the phylogenetic species concept (PSC) have favoured elevations of subspecies to the full species status non-uniformly across the primate tree. As a result, relying on diversification estimates that only consider species would result in disproportionally high diversification rate estimates being assigned to lineages where elevations of subspecies to the full species status have been favoured (e.g. strepsirrhines and platyrrhines; Isaac, et al., 2004).

*Structural Proxies of Behavioural Flexibility*

In addition to behavioural measures, we used brain volume measures as a proxy of behavioural flexibility. We note that while brains perform many functions, and the link between volume and function is not well established (e.g. Healy & Rowe, 2007; Logan, et al., 2018), behavioural flexibility measures such as innovation rate, social learning rate and learning performance in the laboratory do correlate with brain volume measures across species (Reader, 2003; Lefebvre, et al., 2004; Reader, et al., 2011). This suggests that primate brain volume measures are useful secondary proxies of behavioural flexibility. While direct measures of behavioural flexibility under standardized testing conditions would be valuable, such measures are not available for the large-scale comparative tests we conduct here. We can however compare the consistency of results across different proxies for behavioural flexibility.

Data on brain size, estimated as endocranial volume (ECV) in $cm^3$ (hereafter 'brain volume'), were obtained from Powell et al. (2017) (a compilation containing data from Isler, et al., 2008 and van Woerden, et al., 2010; 2012; 2014). Powell et al. (2017) calculated species means for endocranial volume (ECV) across males and females for species that were not considered sexually dimorphic. For sexually dimorphic species (size difference > 10%), Powell et al. (2017) used only female measures of ECV to create means. When assigning these values to species and subspecies in our own dataset, one species, *Semnopithecus dussumieri*, was no longer considered to be a valid taxon (Roos, et al., 2014), and was omitted from our dataset.

Data for neocortex, cerebellum and rest-of-brain volume were obtained from the compilations in DeCasien & Higham (2019) and Navarrete et al. (2018). Navarrete et al. (2018) obtained data through measurements of MRI scans. The DeCasien & Higham (2019) compilation includes data from the prior Reader & Macdonald (2003) compilation and contains brain component volumes ($mm^3$) from multiple studies, including data from both MRI and serial section measurements: Stephan et al. (1970); Stephan et al. (1981); Frahm et al. (1984); Stephan et al. (1988); Rilling & Insel (1998); Rilling & Insel (1999); MacLeod et al. (2003); Bush & Allman (2004a); Bush & Allman (2004b); Sherwood et al. (2004); Sherwood et al. (2005); Barger et al. (2007); De Sousa et al. (2010); Barger et al. (2014); Barks et al. (2015); Bauernfeind et al. (2013); Stimpson et al. (2016). Where necessary, DeCasien & Higham (2019) converted brain component masses to volumetric measurements by dividing mass by the density of fresh tissue (1.036 grams per cubic centimeter). DeCasien & Higham (2019) apply the dataset corrections detailed in Reader & Macdonald (2003).

We only included data on neocortex, cerebellum and rest-of-brain volumes from either source when all these measures were made on the same individuals. We were interested in the underlying neural activity that contributes to behavioural flexibility – often attributed to neuron density (see references in Mikhalevich, et al., 2017) – thus we opted to use only neocortical grey matter volume (regions containing neural cell bodies). In some cases volume estimates were reported as means for multiple individuals (n >1 individuals); in order to weight individual entries equally, estimates from each source were multiplied by their respective n values. These values were then summarized across taxa by summing the individual estimates (previously multiplied by n) and then dividing by the total n value for that species/subspecies across studies prior to creating lineage and genus-level estimates. Brain size and behavioural measures of flexibility (e.g. innovation rate) are often summarized at the genus-level (e.g. Riska & Atchley, 1985; Barton, 2006; Deaner, et al., 2007; Lefebvre, 2013). However, there is considerable variation in species' propensities to exhibit flexible behaviour within many primate families and subfamilies (e.g. subfamily Cercopithecinae contains the genus *Macaca* – a highly flexible genus, and the genus *Theropithecus* – a relatively low scoring genus in terms of flexible behaviour and brain size correlates; see Figure S3), thus we opted to not summarize any of our data beyond the genus-level.

We used residuals from ordinary least squares (OLS) log-log regressions to estimate brain volume relative to body mass, and neocortex and cerebellum volume relative to rest of brain volume at the lineage-level. We chose the OLS method over a phylogenetically corrected regression since the residuals obtained from PGLS models for relative brain size were biased with respect to body size. This is illustrated in Figure S2, where lineage-level relative brain volume residuals are plotted against lineage body mass for both OLS and PGLS models. Largerbodied lineages consistently had more positive relative brain size residuals – such that the effect of body size was not being completely removed from residuals in the phylogenetic model. We note that OLS residuals did still share some correlation with body mass (r=-0.130; Figure S2), however, this correlation was considerably weaker than the correlation observed for the PGLS residuals (r=0.316; Figure S2). Given this, we proceeded with OLS residuals – and the phylogenetic signal in these residuals were later accounted for in downstream PGLS analyses. Lineage-level residuals were averaged to get genus-level estimates (see 'Methods' section of main text). We are limited to averaging data among taxa (subspecies and species) at the lineage-level because our phylogenetic imputation of missing brain volume and body mass data relied on a phylogeny, and thus had to be done at the lineage-level. However, we note that our lineages are typically equivalent to the species for which brain data are available. Past studies using comparable primate brain volume residuals (e.g. Reader & Laland, 2002) have relied on much more conservative species taxonomies which are comparable to our 239 species-complexes (i.e. lineages). Thus, averaging brain volume and body mass estimates at the lineage-level before taking the residuals is the equivalent of calculating residuals at the species-level for most comparative studies of primates. Residual analyses may lead to an inflation of Type II error as it is a very conservative method of controlling for body size (Darlington & Smulders, 2001). However, including both variables in a multiple regression as suggested by some (e.g. Freckleton, 2002) caused errors in parameter estimation in the PGLS models due to the very high correlation between brain volume and body mass, resulting in exceptionally large and opposing effect sizes. As a result, we opted to use the more conservative approach based on residual brain volume where subsequent parameter estimates were similarly conservative.

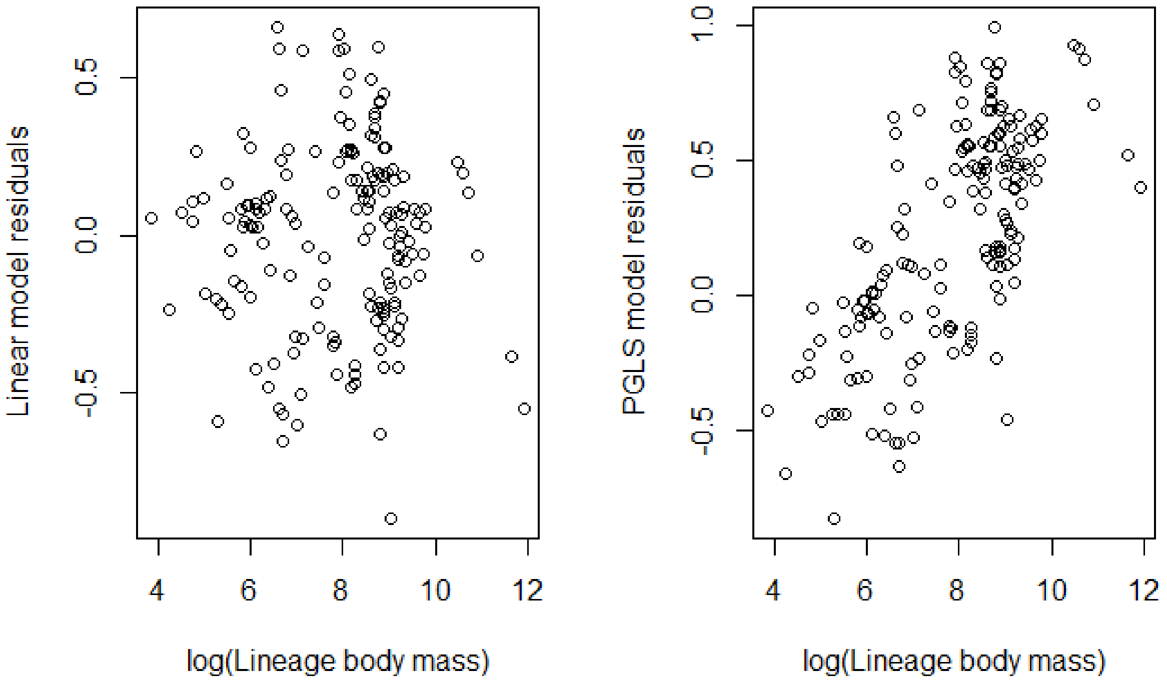


Figure S2: Lineage-level OLS and PGLS residuals versus body mass. Correlation coefficients are r=-0.130 for OLS residuals and r=0.316 for PGLS residuals.

*Behavioural Proxies of Behavioural Flexibility*

We matched behavioural data from Reader et al. (2011) to our own species and subspecies list provided by the IUCN/SSC Primate Specialist Group (2018). One species used by Reader et al. (2011), *Aotus hershkovitzi*, has since been reclassified as a junior synonym of species *Aotus lemurinus* (Defler & Bueno, 2007), and was thus omitted from our analysis. Technical innovation data for species used by Reader et al. (2011), but not Navarrete et al. (2016), were supplemented by examining and categorizing innovation reports provided by Reader et al. (2011). Reader et al. (2011) additionally provide a measure of research effort for each species, recorded as the number of published articles per species published in a survey of the Zoological Record. Some species in the Reader et al. (2011) dataset were reported to have zero accounts of technical innovation and social learning with no recorded measure of research effort or a research effort of zero papers. Data from these species were also omitted from our dataset. Research effort from Reader et al. (2011) was also summarized at both the lineage and genus-levels as a sum of the total number of published articles recorded for each species or subspecies assigned to the given lineage or genus to inform imputation models on the reliability of our observed trait values and to test possible data collection biases associated with research effort.

Experimental data on behavioural innovations and social learning would be preferred to observational data from the literature (Reader & Biro, 2010), but these data are not available for the wide taxonomic spread of our study. The innovation and social learning accounts from literature sources provide quantitative comparative data for a large number of species, typically from observations in the wild. Despite acknowledged weaknesses of such observational data (discussed in Lefebvre, et al., 1997; Laland & Reader, 1999; Reader, 2003; Reader & MacDonald, 2003; Reader, et al., 2011), the taxonomic spread allows for tests of large-scale macroevolutionary trends that can then be followed up by targeted experimental approaches.

*Correlation of Predictors*

Correlation between predictor variables varied greatly (Table S1). Powell et al. (2017) provide their own body mass data from previous compilations, which were highly correlated with body mass estimates from PanTHERIA (Jones, et al., 2009; Table S1). Therefore, we chose to use the more extensive body mass data provided by PanTHERIA in analyses.

*Phylogenetic Patterns*

To visualize phylogenetic patterns in our measures of innovation, social learning, brain size, and diversification rate we painted our 10kTrees consensus tree with these data using the "plotBranchbyTrait" function in the R package phytools (Revell, 2012; 2014), which implements an ancestral-state reconstruction based on symmetric models under maximum likelihood. We also used geiger (Harmon, et al., 2008) to estimate the symmetrical transition rate for the gain and loss of innovation and social learning at the lineage-level, needed to produce the null (simulated) distribution of lineages with innovation and lineages with social learning. We also estimated the phylogenetic signal for all variables of interest. For continuous variables phylogenetic signal was estimated using Pagel's $\lambda$ (Freckleton, et al., 2002), where a $\lambda$ value nearing 1 denotes stronger signal, using the “phylosig” function in phytools. For discrete variables, we used Fritz and Purvis's (2010) D statistic, where a D approaching 0 denotes stronger signal. Results are in Table S2.

*Distribution of Behavioural Flexibility Measures*

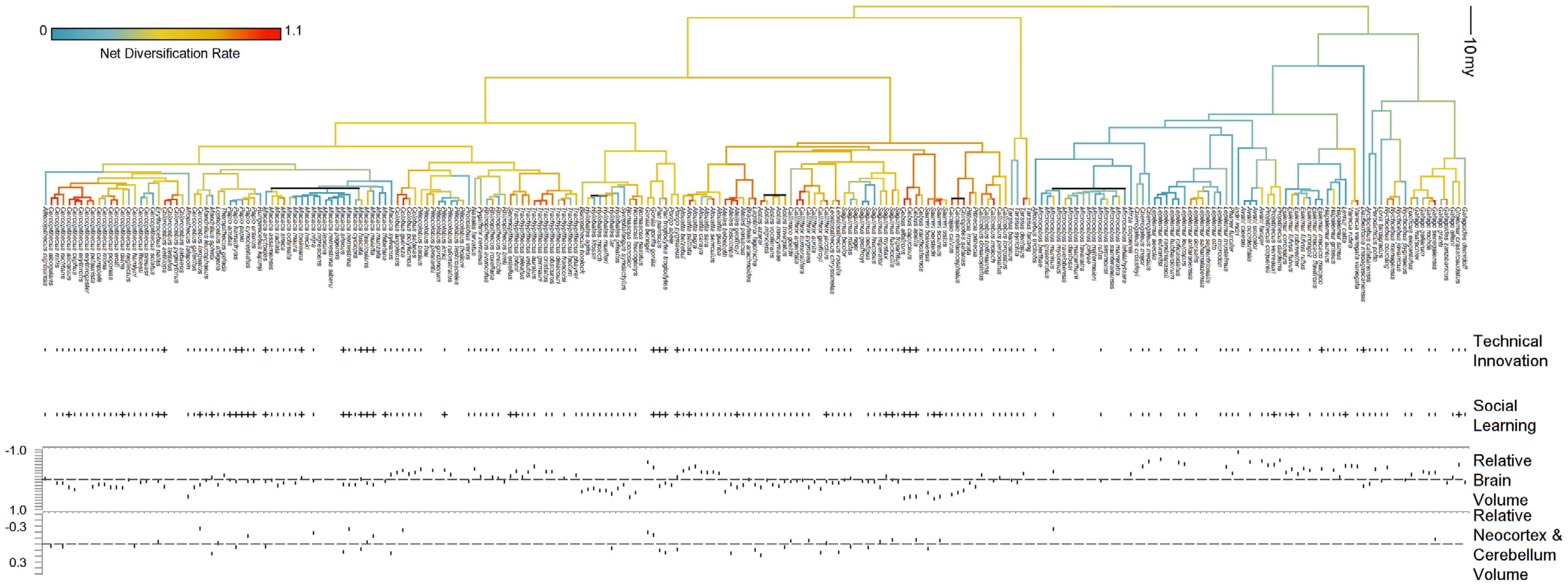


<u>Figure S3:</u> Lineage-level phylogeny painted by Taxa per Lineage Diversification Rate with qualifiers of proxies for behavioural flexibility among non-imputed data. For binary variables (technical innovation and social learning) (+) indicates presence and (–) indicates absence.

*Phylogenetic Imputations of Behavioural Flexibility Measures*

For many of the lineages in our dataset, we were missing data for our different measures and co-variates. Brain data and body mass data are not available for some lineages, while technical innovation and social learning data were collected under a different taxonomy than the one used in this study, meaning that some lineages were not represented in the search done by Reader et al. (2011) due to changes in naming conventions. Additionally, some species searched for by Reader et al. (2011) had no published papers, in which case we considered them to be missing data points. The proportion of missing data varied greatly among variables with body mass (n=196; 82.0%) and relative brain volume (n=186; 77.8%) having the most available data, technical innovation (n=176; 73.6%) and social learning (n=176; 73.6%) having fewer data, and our measures of neocortex, cerebellum and brain volume (n=42; 17.6%) having the least available data (Table S3). Many of the predictors that we examined exhibited strong phylogenetic patterns (Table S2), allowing us to use phylogenetically-informed prediction to estimate the missing traits for these lineages (Pagel, 1999; Guénard, et al., 2013; Swenson, 2014; Penone, et al., 2014). We created a phylogenetic variance-covariance matrix of our pruned 10kTrees phylogeny using the MCMCglmm package (Hadfield, 2010) in R. We decomposed this phylogenetic variance-covariance matrix into 238 eigenvectors using the "PVRdecomp" function from the R package PVR (Santos, et al., 2018), where each eigenvector represents a node in the phylogeny following the order of bifurcation from root to tips. We built phylogenetic models for each proxy of behavioural flexibility (relative brain volume, technical innovation and social learning), using forward model selection to determine the phylogenetic eigenvectors (and covariates) that had the best support for inclusion based on Akaike Information Criterion ($AIC_c$) scores. We implemented this stepwise model selection for phylogenetic models of each covariate using the "stepAIC" function from the MASS R package (Venables & Ripley, 2002).

Our goal for phylogenetic imputation was to maximize the predictive accuracy in our models using phylogenetic position and other covariates to estimate missing data. We ordered our imputation of traits so that we could use imputed information to inform subsequent model fits – for example, body mass is required to estimate brain size. Although there may be concern that this introduces circularity to the analyses, the correlation structure of these covariates is already present in the observed data (Table S1) and imputing with these covariates merely propagates the already present correlations. We first performed model selection on the missing body masses (18.2% of the data) using OLS regression (on the natural log of body mass) of the first 50 phylogenetic eigenvectors (corresponding to the 50 deepest splits in the tree). We then used the phylogenetic eigenvectors with logarithmic body mass as a covariate to estimate brain volume (natural log transformed) with an OLS regression. We used generalized linear models, with a Bernoulli error distribution, for the presence (1) or absence (0) of technical innovation and social learning, with observations weighted by the square-root of research effort. Weighting by research effort, i.e. the number of published articles per species published in the Zoological Record documented by Reader et al. (2011), informs the model about the strength of evidence for a given lineage exhibiting either of these measures of behavioural flexibility: higher research effort indicates a higher certainty that a lineage does or does not display a given behaviour. Innovation and social learning are moderately correlated ($r = 0.533$; Table S1), so we first modelled technical innovation and then used this as a predictive covariate for social learning. The coefficients used in each model can be found in Table S3. We used a larger number of eigenvectors (50; see Table S3) to impute data for variables with high phylogenetic signal and less missing data (e.g. body mass and brain volume) to potentially capture variation presenting itself in shallower splits in the phylogeny. In cases where phylogenetic signal was lower (e.g. technical innovation and social learning) and we had less observed data, we opted to use a lesser number of eigenvectors (40; see Table S3) to avoid problems of overfitting.

We performed a leave-one-out cross-validation to assess the performance of all models. This procedure involved serially removing each lineage's data and using the remaining dataset to replicate our same model-building procedure and predicting that now "missing" lineage's data from the best-fit model. The resulting estimate can then be compared to each lineage's observed value to evaluate how closely we are able to recreate observed data from the models. We evaluated this for continuous variables (body mass and brain volume) based on each model's predictive accuracy ($p^2$):

Equation S1:

$$p^2 = 1 - \sqrt{\frac{\text{mean}(\hat{y} - y)}{\text{variance}(y)}}$$

where ŷ is the predicted value and y is the observed trait value (Greenberg, et al., 2017). Predictive accuracy was high (approaching $p^2=1$) in all cases (Table S3). For our binary variables, technical innovation and social learning, we instead used an approach that quantified whether models could discriminate the presence or absence of these behaviours based on phylogenetic position and covariates. To do this, we constructed receiver operating characteristic curves that compare the true positive and false positive rate across different model predicted probability thresholds for classifying the presence or absence of each behaviour in a species. An accurate model will have a high rate of true positives and negatives, relative to false positives and negatives. The overall model performance (across thresholds) can be assessed by taking the integral of the receiver operating curve: the area under the curve (AUC). The AUC was taken as a measure of classifier performance for binary data. The AUC describes how accurately a given model can predict binary outcomes, and is scored between 1 (describing a perfect fit; model predictions are all correct) and 0 (model predictions are all incorrect), with a score of 0.5 describing chance performance (Bradley, 1997). Receiver operating characteristic curves and AUC values were calculated using the ROCR package (Sing, et al., 2005) (Figure S4). For these binary traits, the AUC was moderate to high (Table S3), indicating good model performance. For prediction of missing data, we had to choose a predicted threshold as a cut-off for binary classification of each behaviour, we chose this threshold by comparing the accuracy (the ratio of true positives + true negatives: false positives + false negatives) of each probability threshold value in the receiver operating characteristic curve and selecting the cut-off associated with optimal accuracy. The genus *Tarsius* consisted of two pairs of sister lineages that were very distantly related to all other primates. Due to their phylogenetic distance from all other lineages it is expected that phylogenetic imputation will perform poorly. As the missing data was generally clustered in species with close relatives we removed genus *Tarsius* from our cross-validation tests to obtain a more representative estimate of model performance for the species that were missing data.

*Lineage-Richness Sampling Bias*

To test whether lineage-rich genera are more likely by chance to have reports of technical innovation or social learning, and to quantify this sampling effect, we simulated the evolution of these traits on our tree independently of diversification rate and tested whether observed correlations depart from this expectation. We simulated the evolution of lineage-specific technical innovation and social learning on the phylogeny over 1000 iterations using the symmetrical rate Mk model of discrete trait evolution (Lewis, 2001) implemented through the geiger package (Harmon, et al., 2008). We parameterized the transition rates for the simulation with the Mk model based on the observed presence and absence of both behaviours across the primate phylogeny. We included imputed data for estimating transition rates of these behaviours, noting that removal of these points did not meaningfully influence estimated transition rates. After simulating the random evolution of technical innovation and social learning across the primate phylogeny 1000 times, we repeated genus-level PGLS analyses over the resulting datasets. From these analyses on the simulated datasets, we obtained a distribution of null effect sizes and compared our estimated effect sizes from the real data to calculate the probability of randomly observing an effect size of that magnitude.

*Research Effort Bias*

Research effort varies considerably across primate lineages, and there is the potential that unknown flexible behaviours exist in currently data-limited species. If well-studied lineages are more likely to have reports of technical innovation or social learning then it is possible that genera containing a greater number of lineages could be more likely to include a well-studied lineage by chance. If true, this could create a bias towards these diverse genera displaying technical innovation or social learning and in turn erroneously suggest that the presence of these behaviours increase diversification rate. Though our data did not suggest a bias toward clades with higher taxonomic richness having lineages with higher research effort (Figures S5 and S6), we also wanted to take research effort into account in our analysis. Previous studies have controlled the total number of innovation or social learning instances recorded for a given lineage by research effort, however, this was not possible in our case (for rationale, see Methods section of main text) and we thus used binary data. When testing binary measures of innovation against diversification rate, others have accounted for research effort by including it as a covariate in the model (Ducatez, et al., 2020). However, we found a similar approach problematic for primates. Large-brained and innovative primates are extensively studied due to their complex behaviour and, in some cases, perhaps of their close relationship to humans (e.g. primates that exhibit habitual tool use are studied to elucidate the origins of human and animal material culture; McGrew, 1992; Visalberghi, 1994; van Schaik, et al., 1999; van Schaik, et al., 2003; Koops, et al., 2014). Thus, some part of research effort on a given species may be driven by such work, making it difficult to disentangle research effort and behavioural flexibility measures. Inspection of residuals from models including research effort as a covariate alongside our binary behavioural measures additionally revealed that this approach resulted in numerous irregularities in terms of which lineages are considered better innovators or social learners given a typical level of research effort (i.e. positive residuals). Well-studied primates such as chimpanzees have hundreds of reports of innovation and social learning, as well as hundreds of publications in the Zoological Record survey used to estimate research effort. The binary measure effectively caps these behaviours at 1, meaning that accounting for research effort with a binary technical innovation or social learning measure will unfairly penalize well-studied but 'truly' innovative species, making it appear that their propensity to innovate is falsely low. Therefore, we instead extended our simulation to consider whether a bias towards having better studied lineages in diverse genera could drive a positive association between behavioural proxies of flexibility and diversification rate by chance.

In this second simulation we "evolved" research effort onto the phylogeny over 1000 iterations in tandem, but independently, of the evolution of technical innovation or social learning. We assume research effort (on the log-scale, bounding it above zero) would change across the tree according to a Brownian motion model, with the rate of change ($\sigma$) and ancestral state being estimated from the distribution of the natural log of research effort in our lineage dataset. Of course, the research effort directed by scientists is not a trait that evolves, but certainly there are a number of lineage characteristics and traits that are likely to attract research effort (e.g. geographic location, diurnality, social group size and behaviour) and we can expect many of these characteristics to be shaped by common descent and share phylogenetic inertia. Based on our empirical dataset, we defined a minimum threshold of 8 studies (or log(research effort) = 2.07) to observe social learning or technical innovation, based on the least-studied species with a record of technical innovation (as it contained the higher threshold; Figure S7). In our simulation we then changed evolved data points for any lineages that had evolved the presence of social learning and technical innovation, turning these lineages into non-innovators, if that lineage's independently simulated research effort was below 8 studies. With these lineages assigned 'hidden states', we then repeated our genus-level analyses testing behavioural proxies of behavioural flexibility against diversification rate over these simulated datasets to get a distribution of expected neutral effect sizes that also include this potential research effort bias. This simulation was designed to mimic the effect false negatives (i.e. understudied innovators or social learners) may have on our results, illustrating whether having better studied lineages in diverse genera by chance could drive a positive association between the presence of technical innovation or social learning and diversification rate independent of biological mechanisms.

SUPPLEMENTARY RESULTS:

*Phylogenetic Patterns*

At the lineage-level relative brain volume showed high phylogenetic signal ($\lambda$=0.989; where $\lambda$ closer to 1 indicates strong phylogenetic signal), with a moderate signal for technical innovation (D=0.187; where D closer to 0 indicates strong phylogenetic signal), a modest signal for social learning (D=0.521), and none for relative neocortex and cerebellum volume ($\lambda$<0.001) as has previously been described when testing other relative measures of neocortex volume (possibly as a result of limited power; Shultz & Dunbar, 2006). Taxa per Lineage Diversification Rate also had a very low phylogenetic signal ($\lambda$=0.179) (see Figure S3; Table S2).

At the genus-level relative brain volume again showed high phylogenetic signal ($\lambda$=0.827). Technical innovation (D=0.438) and social learning (D=0.486) showed modest phylogenetic correlations, while relative neocortex and cerebellum volume showed almost no relationship with phylogeny ($\lambda$<0.001). Taxa per Genus Diversification Rate again had a relatively low phylogenetic signal ($\lambda$=0.293) while Lineage per Genus Diversification Rate showed almost no phylogenetic signal ($\lambda$<0.001) (Table S2).

Table S1: Correlation matrix of predictor variables.

| | Technical innovation (factor) | Social learning (factor) | Body mass[1] | Body mass[2] | Brain volume | Relative brain volume | Neocortex & cerebellum volume | Rest of total brain volume | Relative neocortex & cerebellum volume |
|---|---|---|---|---|---|---|---|---|---|
| **Technical innovation (factor)** | ---- | 0.533 | 0.418 | 0.427 | 0.452 | 0.162 | 0.618 | 0.592 | 0.148 |
| **Social learning (factor)** |  | ---- | 0.337 | 0.343 | 0.381 | 0.113 | 0.481 | 0.484 | -0.127 |
| **Body mass[1]** |  |  | ---- | 0.985 | 0.869 | -0.140 | 0.839 | 0.918 | -0.186 |
| **Body mass[2]** |  |  |  | ---- | 0.916 | -0.122 | 0.880 | 0.942 | -0.149 |
| **Brain volume** |  |  |  |  | ---- | 0.106 | 0.988 | 0.992 | -0.042 |
| **Relative brain volume (residuals)** |  |  |  |  |  | ---- | -0.378 | -0.453 | 0.302 |
| **Neocortex & cerebellum volume** |  |  |  |  |  |  | ---- | 0.979 | 0.024 |
| **Rest of total brain volume** |  |  |  |  |  |  |  | ---- | -0.112 |
| **Relative neocortex & cerebellum volume (residuals)** |  |  |  |  |  |  |  |  | ---- |

Technical innovation and social learning predictors represented by 0/1 integers and coefficients are thus not comparable to coefficients of continuous predictors

Data sources: [1]PanTHERIA; [2]Powell et al., 2017

Relative brain volume data obtained by retaining residuals from a log-log regression of brain volume as a function of body mass[1]

Relative neocortex & cerebellum volume data obtained by retaining residuals from a log-log regression of neocortex & cerebellum volume as a function of rest of total brain volume

Table S2: Phylogenetic signal of predictor variables and diversification rate recorded as Pagel's λ (for continuous variables; where λ closer to 1 indicates signal) or the D-statistic (for binary variables; where D closer to 0 indicates signal).

| Variable (Units) | Lineage-level Pagel's λ or D-statistic | Genus-level Pagel's λ or D-statistic |
|---|---|---|
| **Technical innovation (factor)** | D= 0.187 | D= 0.438 |
| **Social learning (factor)** | D= 0.521 | D= 0.486 |
| **Brain volume ($cm^3$)** | λ= 1.000 | λ= 1.000 |
| **Body mass (g)** | λ= 1.000 | λ= 0.818 |
| **Relative brain volume (residuals)** | λ= 0.989 | λ= 0.827 |
| **Neocortex & cerebellum volume ($mm^3$)** | λ= 1.000 | λ= 1.000 |
| **Rest of brain volume ($mm^3$)** | λ= 1.000 | λ= 1.000 |
| **Relative neocortex & cerebellum volume (residuals)** | λ< 0.001 | λ< 0.001 |
| **Taxa per Lineage Diversification Rate** | λ= 0.179 | ---- |
| **Taxa per Genus Diversification Rate** | ---- | λ= 0.293 |
| **Lineage per Genus Diversification Rate** | ---- | λ< 0.001 |

Table S3: Available data (%), predictive accuracy ($p^2$) and AUC scores for predictors across the lineage-level dataset (239 lineages), with selected coefficients and the number of eigenvectors included in imputation models. N/A: not applicable since imputation was not attempted.

| Variable (Units) | # Lineages with available data (%) | Selected coefficients (including # of eigenvectors selected by model from the phylogeny) | # of eigenvectors | Predictive accuracy ($p^2$) or area under the ROC curve (AUC) |
|---|---|---|---|---|
| **Body mass (g)** | 196 (82.0%) | 10KTrees consensus phylogeny (27 eigenvectors) | 50 | $p^2$=0.999 |
| **ECV (brain volume) ($cm^3$)** | 186 (77.8%) | Body Mass + 10KTrees consensus phylogeny (32 eigenvectors) | 50 | $p^2$=0.990 |
| **Technical innovation (factor)** | 176 (73.6%) | ECV + 10KTrees consensus phylogeny (9 eigenvectors) | 40 | AUC= 0.928 |
| **Social learning (factor)** | 176 (73.6%) | Technical innovation + ECV + 10KTrees consensus phylogeny (21 eigenvectors) | 40 | AUC= 0.699 |
| **Neocortex & cerebellum volume ($mm^3$)** | 42 (17.6%) | N/A | N/A | N/A |
| **Rest of total brain volume ($mm^3$)** | 42 (17.6%) | N/A | N/A | N/A |

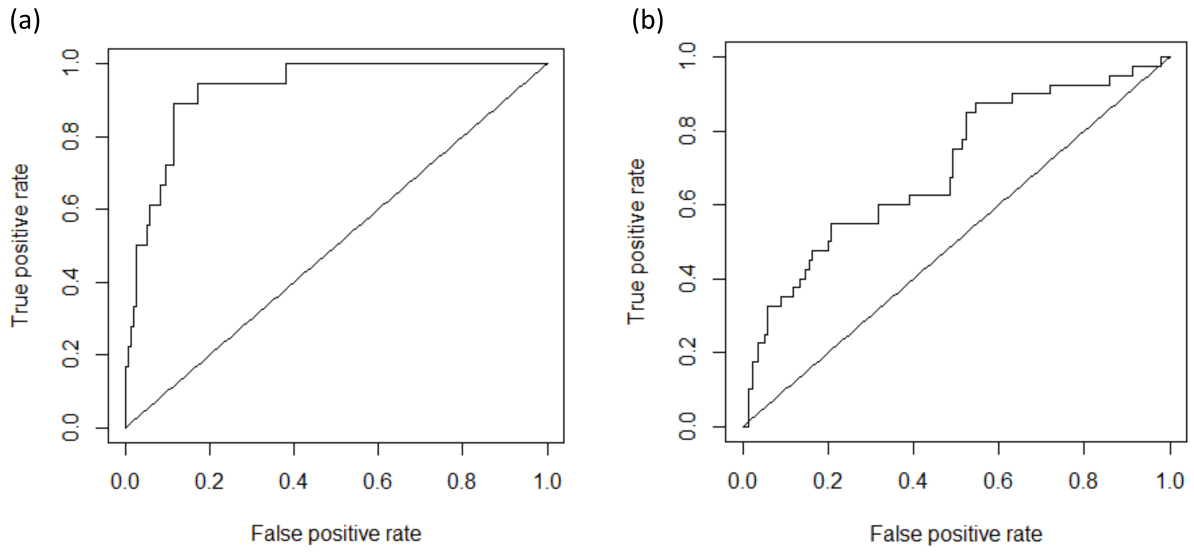


<u>Figure S4:</u> Receiver operating characteristic curves from leave-one-out cross-validation models for predicting the presence or absence of: (a) technical innovation (maximum accuracy=0.926; AUC=0.928) and (b) social learning (maximum accuracy=0.801; AUC=0.699) imputations.

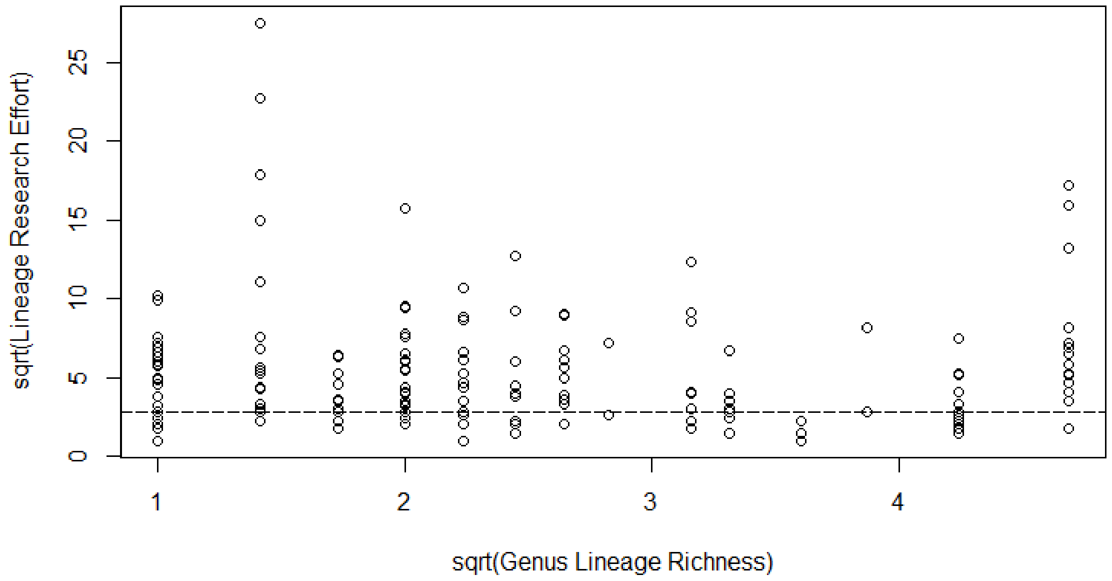


<u>Figure S5:</u> Square-root of lineage per genus richness versus the square-root of lineage-level research effort. The dotted line represents the fewest number of papers in the research effort survey for an observation of technical innovation to be made in a lineage (log(research effort) = 2.07; see Figure S7). There was no trend between lineage research effort and the richness of its genus (ß=-0.239; p=0.343).

(a)

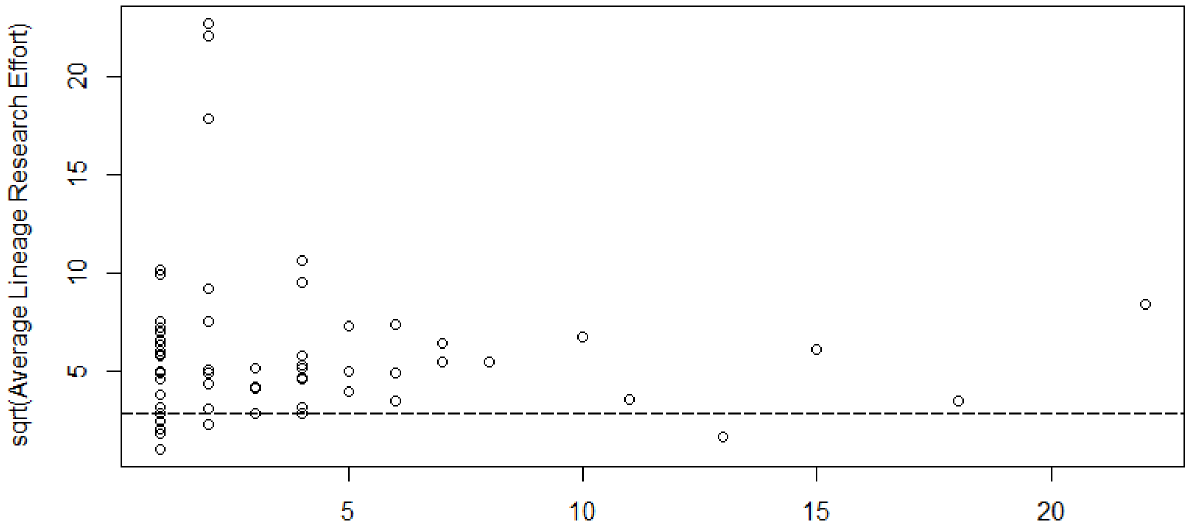


(b)

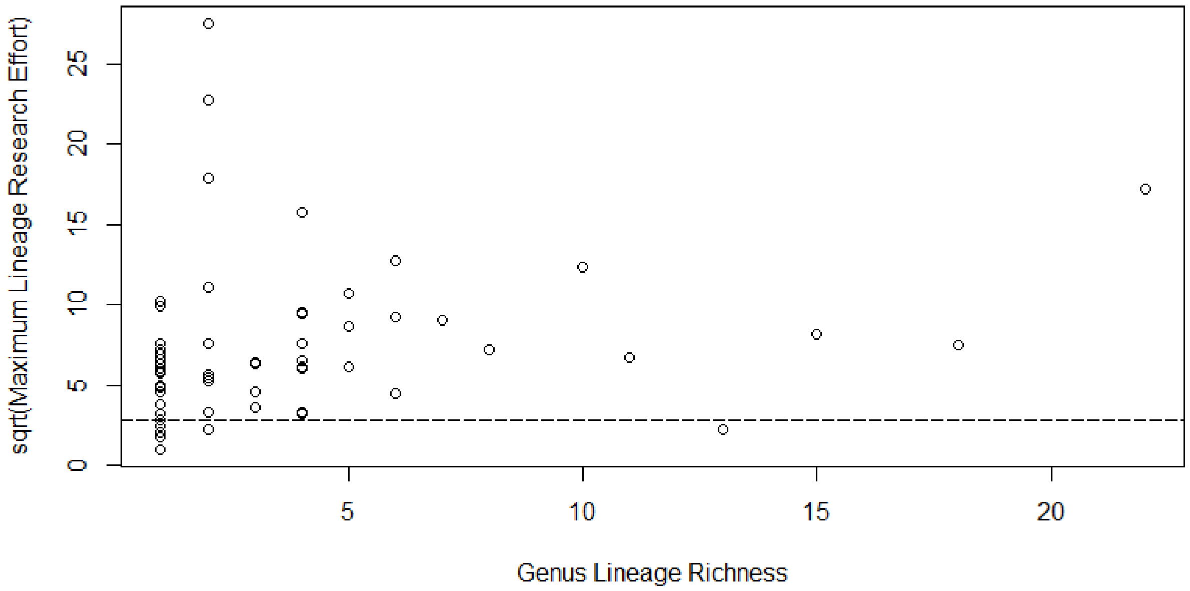


<u>Figure S6:</u> Lineage per genus richness versus (a) the square-root of the average research effort per lineage in each genus, and (b) the square-root of the maximum research effort per lineage in each genus. The dotted line represents the fewest number of papers in the research effort survey for an observation of technical innovation to be made in a lineage (log(research effort) = 2.07; see Figure S7).

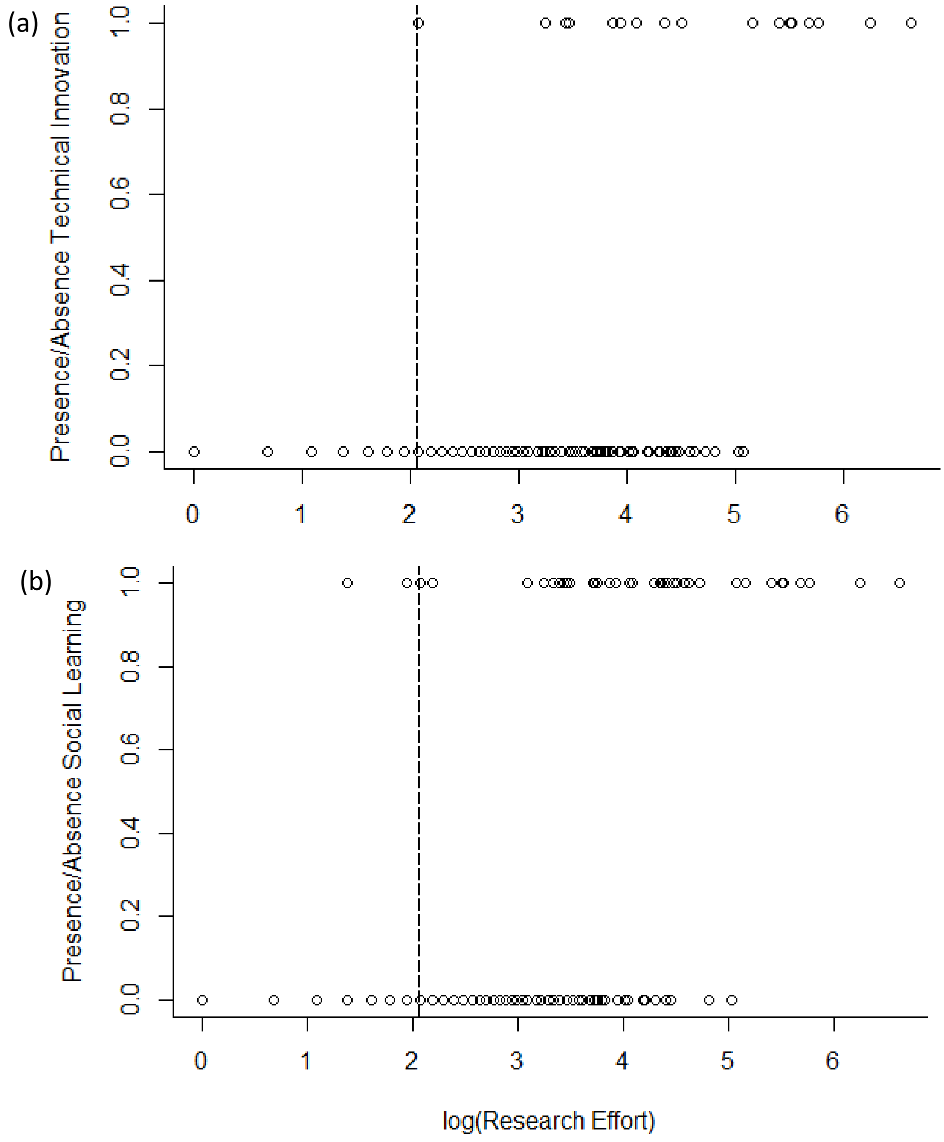


Figure S7: Presence and absence of (a) technical innovation and (b) social learning versus research effort per lineage. The dotted line represents the fewest number of papers in the research effort survey for an observation of technical innovation to be made in a lineage (log(research effort) = 2.07).

Table S4: Results of lineage-level analyses using Taxa per Lineage Diversification Rate from all imputed and non-imputed models +P≤0.1; *P≤0.05; **P≤0.01. Each row represents a separate analysis.

| Variable (Units) | Imputed | | | | | | Non-imputed | | | | | |
|---|---|---|---|---|---|---|---|---|---|---|---|---|
| | Min 95%CI | Max 95%CI | ß | SE | t | p | Min 95%CI | Max 95%CI | ß | SE | t | p |
| **Technical innovation (factor)** | -0.143 | 0.109 | -0.017 | 0.064 | -0.264 | 0.792 | -0.166 | 0.149 | -0.009 | 0.080 | -0.110 | 0.912 |
| **Social learning (factor)** | -0.029 | 0.163 | 0.067 | 0.049 | 1.374 | 0.171 | -0.020 | 0.205 | 0.092 | 0.057 | 1.605 | 0.110 |
| **Technical innovation and social learning (factor)** | -0.133 | 0.130 | -0.002 | 0.067 | -0.026 | 0.979 | -0.151 | 0.185 | 0.017 | 0.086 | 0.200 | 0.841 |
| **Relative brain volume*** | -0.036 | 0.159 | 0.062 | 0.050 | 1.243 | 0.215 | -0.007 | 0.218 | 0.106 | 0.057 | 1.844 | 0.067+ |
| **Relative neocortex & cerebellum volume*** | | | | | | | -0.240 | 0.152 | -0.044 | 0.100 | -0.437 | 0.664 |
| **Body mass*** | -0.107 | 0.145 | 0.019 | 0.064 | 0.300 | 0.764 | -0.111 | 0.147 | 0.018 | 0.066 | 0.270 | 0.787 |

* = $log_e$-transformed and scaled by 2 standard deviation

Table S5: Results of genus-level analyses using Taxa per Genus Diversification Rate from all imputed and non-imputed models +P≤0.1; *P≤0.05; **P≤0.01. Each row represents a separate analysis.

| Variable (Units) | Imputed | | | | | | Non-imputed | | | | | |
|---|---|---|---|---|---|---|---|---|---|---|---|---|
| | Min 95%CI | Max 95%CI | ß | SE | t | p | Min 95%CI | Max 95%CI | ß | SE | t | p |
| **Technical innovation (factor)** | -0.089 | 0.160 | 0.036 | 0.064 | 0.562 | 0.577 | -0.093 | 0.154 | 0.031 | 0.063 | 0.486 | 0.629 |
| **Social learning (factor)** | -0.057 | 0.130 | 0.037 | 0.048 | 0.774 | 0.442 | -0.065 | 0.121 | 0.028 | 0.048 | 0.589 | 0.558 |
| **Technical innovation and social learning (factor)** | -0.075 | 0.185 | 0.055 | 0.066 | 0.828 | 0.411 | -0.081 | 0.177 | 0.048 | 0.066 | 0.733 | 0.467 |
| **Relative brain volume*** | -0.017 | 0.183 | 0.083 | 0.051 | 1.635 | 0.108 | 0.049 | 0.228 | 0.138 | 0.046 | 3.020 | 0.004** |
| **Relative neocortex & cerebellum volume*** | | | | | | | -0.141 | 0.154 | 0.007 | 0.075 | 0.089 | 0.930 |
| **Body mass*** | -0.039 | 0.186 | 0.074 | 0.057 | 1.285 | 0.204 | -0.034 | 0.194 | 0.080 | 0.058 | 1.379 | 0.173 |

* = $log_e$-transformed and scaled by 2 standard deviation

Table S6: Results of genus-level analyses using Lineage per Genus Diversification Rate from all imputed and non-imputed models +P≤0.1; *P≤0.05; **P≤0.01. Each row represents a separate analysis.

| Variable (Units) | Imputed | | | | | | Non-imputed | | | | | |
|---|---|---|---|---|---|---|---|---|---|---|---|---|
| | Min 95%CI | Max 95%CI | ß | SE | t | p | Min 95%CI | Max 95%CI | ß | SE | t | p |
| **Technical innovation (factor)** | 0.002 | 0.130 | 0.066 | 0.033 | 2.025 | 0.048* | 0.000 | 0.129 | 0.065 | 0.033 | 1.973 | 0.053+ |
| **Social learning (factor)** | 0.043 | 0.132 | 0.088 | 0.023 | 3.884 | < 0.001** | 0.042 | 0.131 | 0.086 | 0.023 | 3.789 | < 0.001** |
| **Technical innovation and social learning (factor)** | 0.018 | 0.151 | 0.085 | 0.034 | 2.504 | 0.015* | 0.017 | 0.150 | 0.083 | 0.034 | 2.452 | 0.017* |
| **Relative brain volume*** | -0.022 | 0.072 | 0.025 | 0.024 | 1.037 | 0.304 | -0.019 | 0.078 | 0.030 | 0.025 | 1.203 | 0.234 |
| **Relative neocortex & cerebellum volume*** | | | | | | | -0.112 | 0.074 | -0.019 | 0.047 | -0.400 | 0.693 |
| **Body mass*** | -0.020 | 0.074 | 0.027 | 0.024 | 1.118 | 0.268 | -0.017 | 0.078 | 0.030 | 0.024 | 1.247 | 0.218 |

* = $\log_e$-transformed and scaled by 2 standard deviation

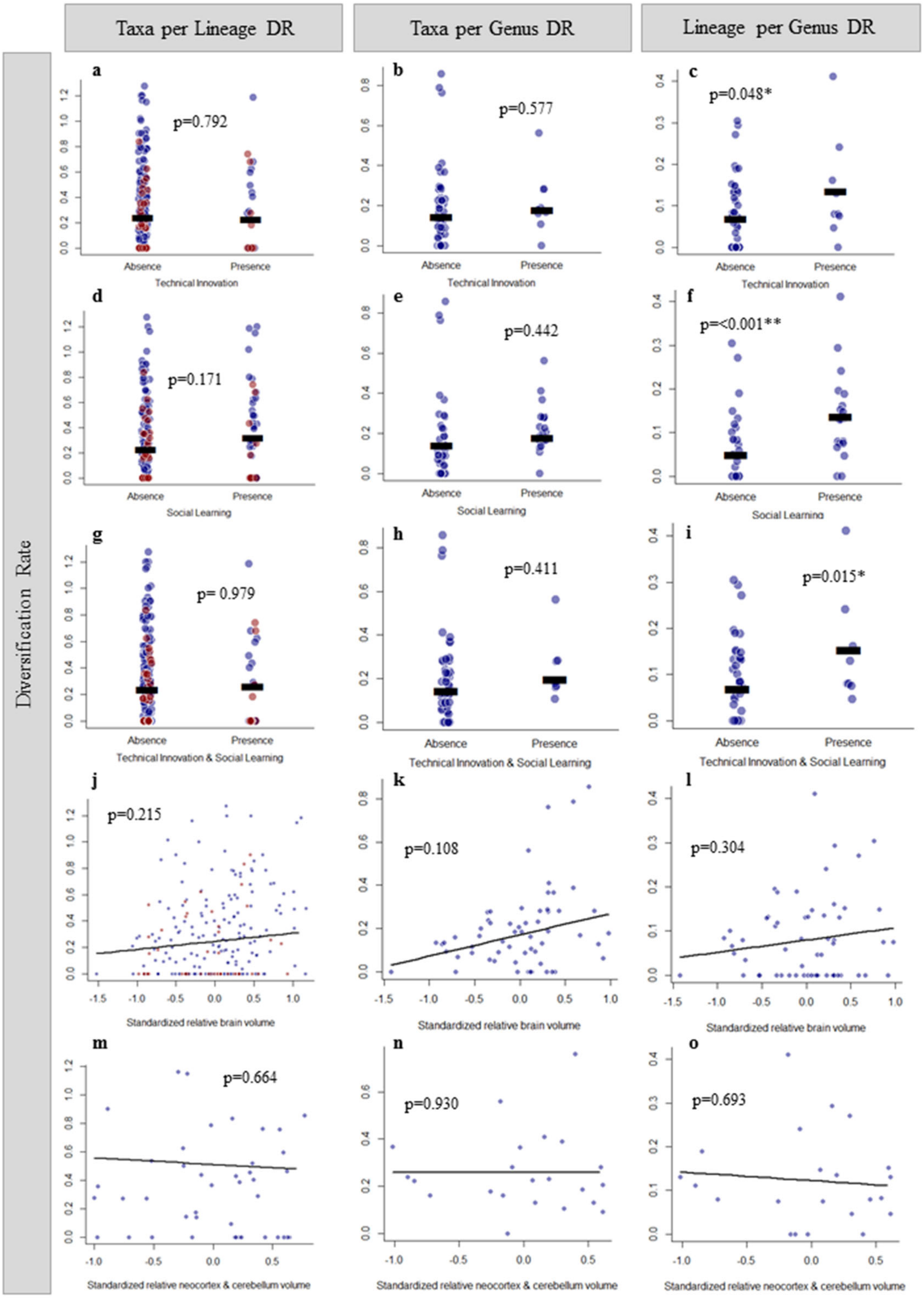


Figure S8: Relationships between technical innovation (a-c), social learning (d-f), combined technical innovation and social learning (g-i), relative brain volume (j-l) (all including imputed data) and relative neocortex & cerebellum volume (m-o) (non-imputed) on three measures of primate diversification rate. Imputed lineage-level data points are indicated in red. Horizontal bars indicate group means. Significance indicated as: +P≤0.1; *P≤0.05; **P≤0.01.

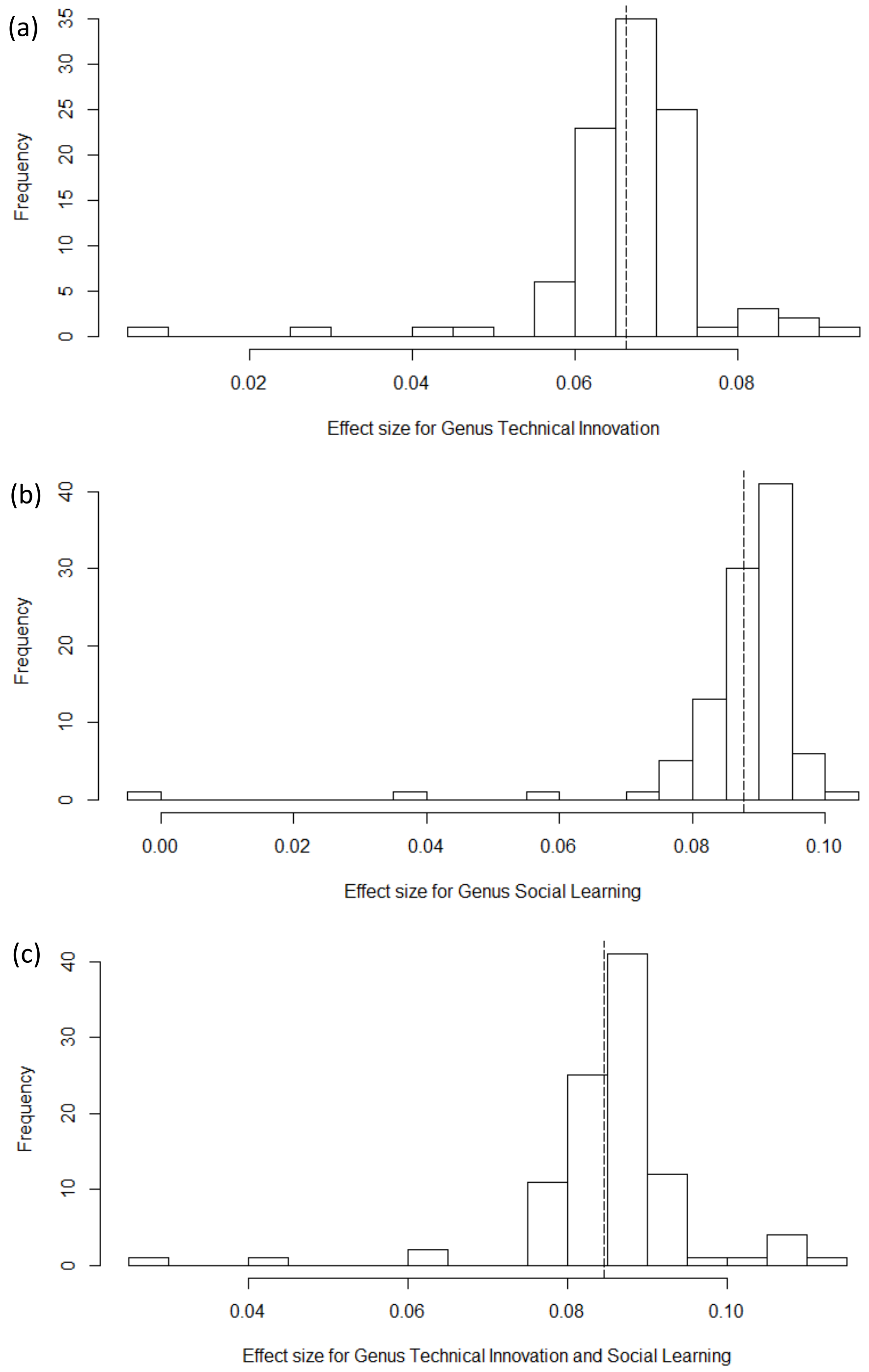


Figure S9: Distribution of effect sizes from 100 phylogenetic generalized least squares (PGLS) regressions testing the relationship between Lineage per Genus Diversification Rate and (a) technical innovation (b) social learning, and (c) the combined presence of both using a tree block of 1000 randomly sampled trees from 10kTrees (Arnold, et al., 2010). The dotted line indicates the observed effect size reported in Table S6.

*Lineage-Richness Sampling Bias*

From our simulations, we confirmed that there is a bias towards a positive relationship between diversification rate and the presence of flexible behaviours (Figure S10). For Taxa per Genus Diversification Rate, the probabilities of the observed effect sizes (ß) occurring by chance were $p=0.392$ for presence of technical innovation, $p=0.420$ for presence of social learning, and $p=0.260$ for combined presence of technical innovation and social learning. For Lineage per Genus Diversification Rate, the probabilities of the observed effect sizes (ß) occurring by chance were $p=0.028$ for presence of technical innovation, $p=0.002$ for presence of social learning, and $p=0.008$ for combined presence of technical innovation and social learning. That is, the effect sizes generated by this bias for the association with Lineage per Genus Diversification Rate were considerably lower than the observed effect size from the empirical data.

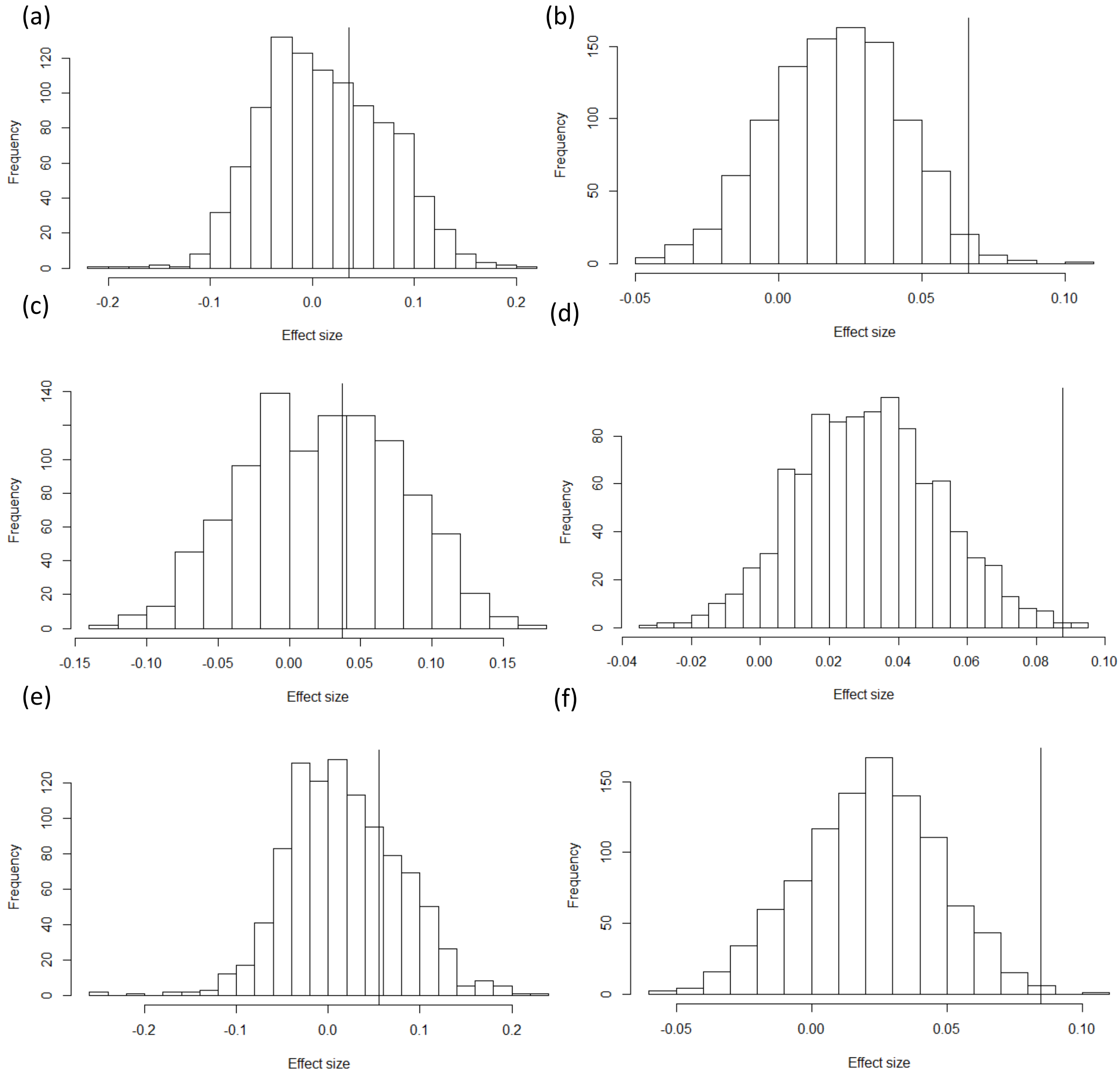


Figure S10: Distribution of effect sizes (ß) from PGLS tests on 1000 simulations of the random evolution of technical innovation and social learning. The neutral effect sizes for technical innovation on (a) Taxa per Genus Diversification Rate (DR) (median ß=0.009) and (b) Lineage per Genus DR (median ß=0.020); effect sizes for social learning on (c) Taxa per Genus DR (median ß=0.025) and (d) Lineage per Genus DR (median ß=0.031); Effect sizes for the combined presence of technical innovation and social learning on (e) Taxa per Genus DR (median ß=0.012) and (f) Lineage per Genus DR (median ß=0.023). Vertical line indicates the observed effect size for each predictor.

*Research Effort Bias*

Positive correlations between our measures of behavioural flexibility and genus-level estimates of diversification rate could also be further biased by asymmetry in research effort. If many understudied lineages scored as zero are truly technical innovators or social learners (i.e. false negatives), and these lineages tend to be in depauperate clades, then heightened observation of behavioural flexibility in diverse clades could bias our results. Testing this potential bias with a simulation where observed behaviours were hidden based on simulated research effort, we found that this scenario generated a slightly greater bias towards a positive association between diversification rate and the presence of each behavioural flexibility measure (Figure S11). Our results for Taxa per Genus Diversification Rate remained non-significant, and for Lineage per Genus Diversification Rate remained significant except for technical innovation. For Taxa per Genus Diversification Rate, the probabilities of the observed effect sizes (ß) occurring by chance were p=0.408 for presence of technical innovation, p=0.439 for presence of social learning, and p=0.281 for combined presence of technical innovation and social learning. For Lineage per Genus Diversification Rate, the probabilities of the observed effect sizes (ß) occurring by chance were p=0.109 for presence of technical innovation, p=0.019 for presence of social learning, and p=0.027 for combined presence of technical innovation and social learning.

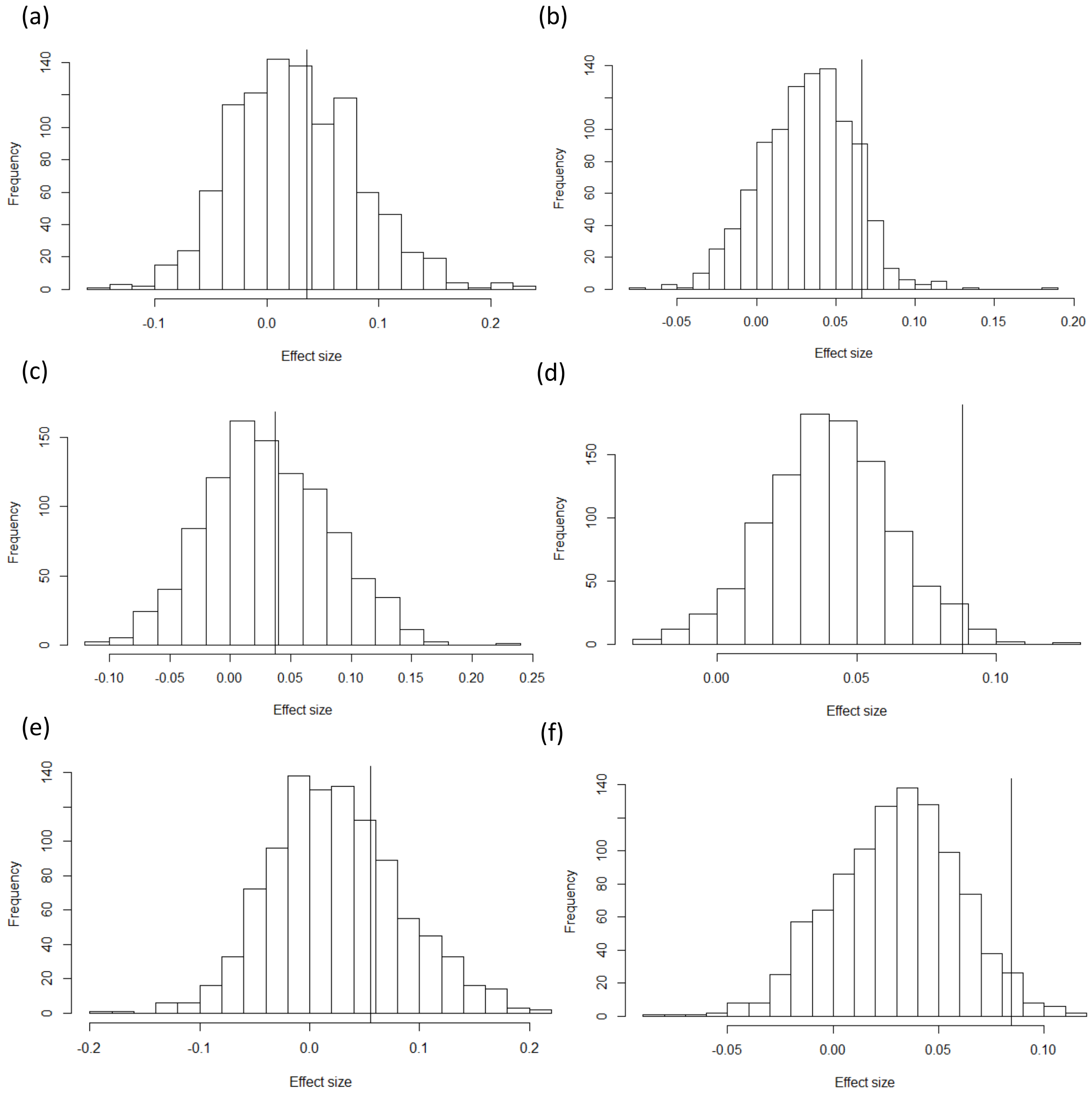


Figure S11: Null distribution of effect sizes (ß) from PGLS tests on 1000 simulated technical innovation/social learning datasets censoring the presence of these behaviours when a lineage has a simulated research effort < 8 papers, testing the effect of technical innovation on (a) Taxa per Genus Diversification Rate (DR) (median ß=0.023) and (b) Lineage per Genus DR (median ß=0.033); social learning on (c) Taxa per Genus DR (median ß=0.028) and (d) Lineage per Genus DR (median ß=0.040); and the combined presence of technical innovation and social learning on (e) Taxa per Genus DR (median ß=0.020) and (f) Lineage per Genus DR (median ß=0.032). Vertical line indicates the observed effect size.

Table S7: Results of lineage-level analyses after dropping great apes using Taxa per Lineage Diversification Rate from all imputed and non-imputed models +P≤0.1; *P≤0.05; **P≤0.01. Each row represents a separate analysis.

| Variable (Units) | Imputed | | | | | | Non-imputed | | | | | |
|---|---|---|---|---|---|---|---|---|---|---|---|---|
| | Min 95%CI | Max 95%CI | ß | SE | t | p | Min 95%CI | Max 95%CI | ß | SE | t | p |
| **Technical innovation (factor)** | -0.160 | 0.120 | -0.020 | 0.071 | -0.276 | 0.783 | -0.187 | 0.165 | -0.011 | 0.090 | -0.123 | 0.903 |
| **Social learning (factor)** | -0.027 | 0.175 | 0.074 | 0.051 | 1.440 | 0.151 | -0.018 | 0.218 | 0.100 | 0.060 | 1.665 | 0.098+ |
| **Technical innovation and social learning (factor)** | -0.149 | 0.147 | -0.001 | 0.076 | -0.016 | 0.988 | -0.169 | 0.212 | 0.022 | 0.097 | 0.221 | 0.825 |
| **Relative brain volume*** | -0.034 | 0.170 | 0.068 | 0.052 | 1.308 | 0.192 | -0.004 | 0.230 | 0.113 | 0.060 | 1.890 | 0.061+ |
| **Relative neocortex & cerebellum volume*** | | | | | | | -0.280 | 0.149 | -0.066 | 0.109 | -0.598 | 0.553 |
| **Body mass*** | -0.111 | 0.152 | 0.020 | 0.067 | 0.301 | 0.763 | -0.113 | 0.155 | 0.021 | 0.068 | 0.303 | 0.762 |

* = $\log_e$-transformed and scaled by 2 standard deviation

Table S8: Results of genus-level analyses after dropping great apes using Taxa per Genus Diversification Rate from all imputed and non-imputed models +P≤0.1; *P≤0.05; **P≤0.01. Each row represents a separate analysis.

| Variable (Units) | Imputed | | | | | | Non-imputed | | | | | |
|---|---|---|---|---|---|---|---|---|---|---|---|---|
| | Min 95%CI | Max 95%CI | ß | SE | t | p | Min 95%CI | Max 95%CI | ß | SE | t | p |
| **Technical innovation (factor)** | -0.076 | 0.219 | 0.072 | 0.075 | 0.953 | 0.345 | -0.081 | 0.211 | 0.065 | 0.074 | 0.870 | 0.388 |
| **Social learning (factor)** | -0.051 | 0.149 | 0.049 | 0.051 | 0.960 | 0.341 | -0.060 | 0.140 | 0.040 | 0.051 | 0.778 | 0.440 |
| **Technical innovation and social learning (factor)** | -0.053 | 0.262 | 0.105 | 0.080 | 1.304 | 0.198 | -0.061 | 0.250 | 0.095 | 0.079 | 1.195 | 0.237 |
| **Relative brain volume*** | -0.018 | 0.194 | 0.088 | 0.054 | 1.634 | 0.108 | 0.050 | 0.238 | 0.144 | 0.048 | 2.999 | 0.004** |
| **Relative neocortex & cerebellum volume*** | | | | | | | -0.151 | 0.182 | 0.015 | 0.085 | 0.181 | 0.859 |
| **Body mass*** | -0.015 | 0.213 | 0.099 | 0.058 | 1.700 | 0.095+ | -0.007 | 0.222 | 0.108 | 0.058 | 1.847 | 0.070+ |

* = $\log_e$-transformed and scaled by 2 standard deviation

Table S9: Results of genus-level analyses after dropping great apes using Lineage per Genus Diversification Rate from all imputed and non-imputed models +P≤0.1; *P≤0.05; **P≤0.01. Each row represents a separate analysis.

| **Variable (Units)** | **Imputed** | | | | | | **Non-imputed** | | | | | |
|---|---|---|---|---|---|---|---|---|---|---|---|---|
| | Min 95%CI | Max 95%CI | ß | SE | t | p | Min 95%CI | Max 95%CI | ß | SE | t | p |
| **Technical innovation (factor)** | 0.023 | 0.177 | 0.100 | 0.039 | 2.549 | 0.014* | 0.021 | 0.175 | 0.098 | 0.039 | 2.505 | 0.015* |
| **Social learning (factor)** | 0.052 | 0.146 | 0.099 | 0.024 | 4.144 | < 0.001** | 0.050 | 0.145 | 0.098 | 0.024 | 4.049 | < 0.001** |
| **Technical innovation and social learning (factor)** | 0.054 | 0.216 | 0.135 | 0.041 | 3.285 | 0.002** | 0.053 | 0.215 | 0.134 | 0.041 | 3.238 | 0.002** |
| **Relative brain volume*** | -0.023 | 0.076 | 0.026 | 0.025 | 1.042 | 0.302 | -0.019 | 0.082 | 0.031 | 0.026 | 1.208 | 0.233 |
| **Relative neocortex & cerebellum volume*** | | | | | | | -0.123 | 0.089 | -0.017 | 0.054 | -0.309 | 0.761 |
| **Body mass*** | -0.016 | 0.083 | 0.033 | 0.025 | 1.322 | 0.192 | -0.012 | 0.088 | 0.038 | 0.025 | 1.487 | 0.143 |

* = $log_e$-transformed and scaled by 2 standard deviation
